# Phylogeographic reconstruction of the emergence and spread of Powassan virus in the northeastern United States

**DOI:** 10.1101/2022.10.14.512245

**Authors:** Chantal B.F. Vogels, Doug E. Brackney, Alan P. Dupuis, Rebecca M. Robich, Joseph R. Fauver, Anderson F. Brito, Scott C. Williams, John F. Anderson, Charles B. Lubelczyk, Rachel E. Lange, Melissa A. Prusinski, Laura D. Kramer, Jody L. Gangloff-Kaufmann, Laura B. Goodman, Guy Baele, Robert P. Smith, Philip M. Armstrong, Alexander T. Ciota, Simon Dellicour, Nathan D. Grubaugh

## Abstract

Powassan virus is an emerging tick-borne virus of concern for public health, but very little is known about its transmission patterns and ecology. Here, we expanded the genomic dataset by sequencing 279 Powassan viruses isolated from *Ixodes scapularis* ticks from the northeastern United States. Our phylogeographic reconstructions revealed that Powassan virus lineage II was likely introduced or emerged from a relict population in the Northeast between 1940-1975. Sequences strongly clustered by sampling location, suggesting a highly focal geographical distribution. Our analyses further indicated that Powassan virus lineage II emerged in the northeastern U.S. mostly following a south to north pattern, with a weighted lineage dispersal velocity of ~3 km/year. Since the emergence in the Northeast, we found an overall increase in the effective population size of Powassan virus lineage II, but with growth stagnating during recent years. The cascading effect of population expansion of white-tailed deer and *I. scapularis* populations likely facilitated the emergence of Powassan virus in the northeastern U.S.

**Significance statement:** Our work provides important fundamental insights in the local transmission dynamics of an emerging tick-borne pathogen of public health concern. Without the availability of vaccines or specific treatments, prevention of Powassan virus infection is dependent on education and control. We identified that Powassan virus is maintained in highly localized transmission foci that have been maintained for several years, without introductions of new virus clades. This provides both opportunities for better education about high risk areas and effective targeted control in Powassan virus foci with a long lasting impact.

## Introduction

Reports of tick-borne diseases in the United States have been steadily rising, with more than 50,000 cases in 2019 (1). In that same year, a record number of 43 human cases of infection with an emerging tick-borne pathogen, Powassan virus (*Flaviviridae: Flavivirus*), were reported (2–5). Powassan virus infection can cause severe neuroinvasive disease with long-lasting sequelae and high fatality rates in humans. Since its initial identification in 1958 (6), incidence rates of Powassan neuroinvasive disease in humans have dramatically risen in the United States, particularly during recent years in the Northeast (7). As Powassan virus infection is difficult to clinically diagnose (8) and most infections are asymptomatic (3), the reported cases are likely a vast underestimation of the true burden. The increasing number of human infections coupled with the lack of an effective vaccine or medicines, highlights the need to better understand local virus transmission patterns to guide targeted prevention and control measures.

Emergence of tick-borne diseases such as Lyme borreliosis, anaplasmosis, and babesiosis are associated with the spread of *Ixodes scapularis* ticks following expansion of suitable habitats (9) and the reintroduction of their primary adult stage hosts, white-tailed deer (*Odocoileus virginianus*), into the northeastern United States (10, 11). Powassan virus consists of two genetically distinct lineages, of which lineage II, also thought to be primarily maintained by *I. scapularis* ticks and small mammals (12, 13) followed a similar path (3–5). However, very little is known about its ecology and transmission patterns. Genetic approaches, including phylogenetic and phylogeographic inference, are powerful tools to understand patterns of pathogen transmission and spread. However, until recently there were only 23 near-complete Powassan virus genomes available from the United States. Thus, to inform future control efforts and mitigate public health risks, there is a critical need for new and innovative phylogeographic approaches to uncover how Powassan virus is being maintained in the Northeast and which factors facilitate or prevent its spread to other areas.

To investigate, we partnered with public health laboratories throughout the Northeast to sequence Powassan virus isolated from *I. scapularis* ticks in Connecticut, New York, and Maine. We sequenced samples collected in 2008-2019 from both historic endemic sites and from novel regions on the leading edge of expansion in the Northeast. With this expanded genomic dataset, we performed discrete and continuous phylogeographic analyses to answer important questions on the emergence and spread of Powassan virus lineage II, such as: (**1**) When did Powassan virus lineage II emerge in the Northeast? (**2**) How is Powassan virus lineage II locally maintained? (**3**) What are the patterns and velocity of spread? (**4**) Can increased transmission explain the recent increase in reported human cases? (**5**) What is the impact of environmental factors on the dispersal dynamics? Overall, our study provides important insights in the emergence and spread of Powassan virus in the northeastern U.S. Our findings help to better identify potential high-risk areas for exposure, which will in turn help to direct future control efforts.

## Results

### Powassan virus phylogeny

Powassan virus consists of two genetically and ecologically distinct lineages (lineage I and II; **Fig. 1**). Prior to this study, there were 23 near-complete Powassan virus genome sequences publicly available from the United States. In this study, we sequenced an additional 279 Powassan viruses (2 belonging to lineage I, and 277 belonging to lineage II) from positive tick pools identified by public health laboratories in Connecticut, New York, and Maine from 2008-2019 (**Tables S1–3**). We created Nextstrain pages to visualize the Powassan genomic data with builds for all available genomes (14), and a more specific build for genomes available from the northeastern U.S. (15). Publicly available lineage I sequences were available from Russia, Canada (initial case in North America), and the United States, but Powassan virus lineage I had not been reported from the United States since the late 1970s (16, 17). As part of this study, we sequenced two lineage I genomes detected in *I. scapularis* ticks in 2019 from New York (18). Later, we sequenced two additional lineage I genomes detected in *I. cookei* (2020) and *Dermacentor variabilis* (2021) also from New York (19). Identification of Powassan virus lineage I in various tick species highlights how increased virus genomic surveillance can help to expand our knowledge of virus ecology.

**Fig. 1:**
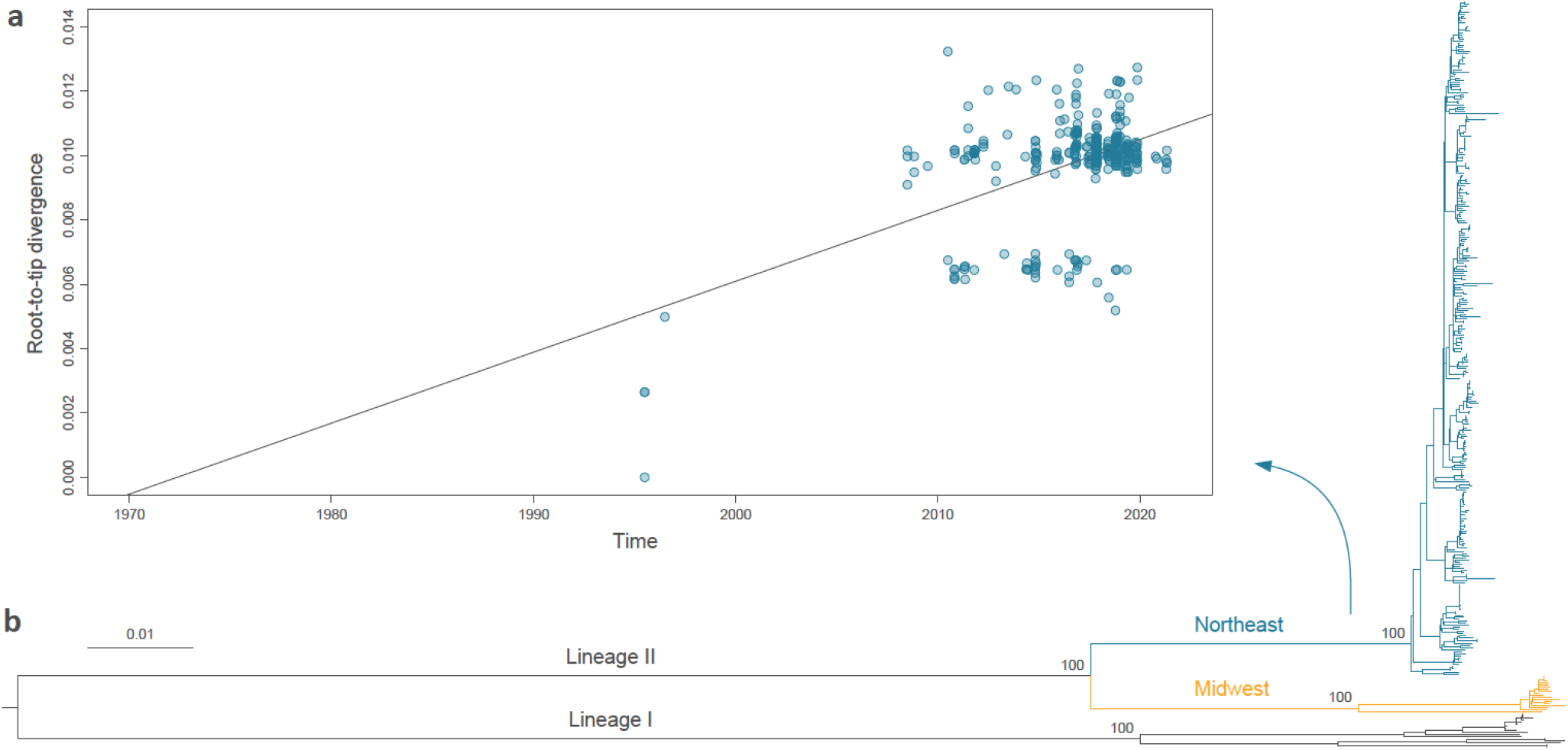
Phylogenetic analysis of Powassan virus lineages. **(a)**Root-to-tip regression performed to assess the temporal signal within the Northeast clade (determination coefficient R2 from the linear regression = 0.23). **(b)**Maximum likelihood tree was obtained from the phylogenetic analysis of publicly available Powassan virus genomes from the United States, Canada, and Russia. Powassan virus lineage II consists of two geographically separated clades in the Northeast and Midwest. Bootstraps support values (based on 1,000 replicates) are provided for the main internal nodes of the tree.

Almost all of the more recent Powassan viruses detected from the US, including what we sequenced for this study, belong to lineage II. Powassan virus lineage II, also referred to as “deer tick virus”, consists of two geographically separated clades comprising viruses from the Midwest and the Northeast (**Fig. 1**). Powassan virus lineage II is the most prevalent in the Northeast, and we first carefully assessed the presence of temporal signal in these data. While the determination coefficient of the root-to-tip regression performed with TempEst (20) is relatively small (R^2^ = 0.23; **Fig. 1**), we find very strong evidence in favor of temporal signal in the data set using a recently developed Bayesian method (21) (log Bayes factor = 41.8 (22)), enabling the use of molecular clocks to estimate time-calibrated phylogenies. As part of this analysis, we find that an uncorrelated relaxed clock with an underlying lognormal distribution provides a better model fit to the data compared to a strict clock model (**Table 1**). We estimate that the evolutionary rate of this clade is 8.25×10^−5^ substitutions/site/year (95% highest posterior density [HPD] interval: [8.23-10.46×10^−5^]; 95% HPD interval of the associated coefficient of variation: [0.48-0.75]). Our estimate is higher than previous estimated evolutionary rates for all Powassan viruses (3.3×10^−5 (23)^) and NS5 coding sequences (3.9×10^−5^-5.4×10^−5 (5, 24)^), but lower than previous estimates based on envelope coding sequences (2.2×10^−4 (5)^). This is likely a reflection of the recent emergence of lineage II in the region. Our work increased the number of publicly available Powassan virus lineage II sequences by more than ten-fold, enabling us to better understand the patterns of emergence and spread in the northeastern U.S.

**Table 1:** Bayesian Evaluation of Temporal Signal analysis.

| **Clock model** | **Model fit** |
| --- | --- |
| Strict clock + dates | -36575.48 |
| Strict clock - dates | -36601.19 |
| Relaxed clock + dates | -36512.57 |
| Relaxed clock - dates | -36554.33 |

### Emergence in the Northeast

The emergence of Powassan virus lineage II into the northeastern U.S. likely followed the re-emergence of *I. scapularis* ticks, its primary vector, into the region. *Ixodes scapularis* originally colonized the Northeast thousands of years ago (25, 26); however, deforestation and restriction of white-tailed deer populations (primary reproductive host for adult *I. scapularis* ticks) during the 1800s greatly reduced *I. scapularis* populations in the Northeast (10, 27). Reforestation and increasing white-tailed deer populations during the mid-1900s then led to a re-emergence of *I. scapularis (10, 28, 29)*.

Our expanded Powassan virus genomic data enabled us to reconstruct the dispersal history of Powassan virus lineage II in the northeastern U.S. (**Fig. 2a**). Our discrete phylogeographic analysis estimates that the time to the most recent common ancestor (tMRCA) for the Northeast Powassan virus lineage II clade is between 1940.3-1974.7 (95% HPD interval; mean 1957.9). This means that lineage II emerged in the Northeast likely before the mid 1970s, corresponding to the re-emergence of *I. scapularis* ticks.

**Fig. 2:**
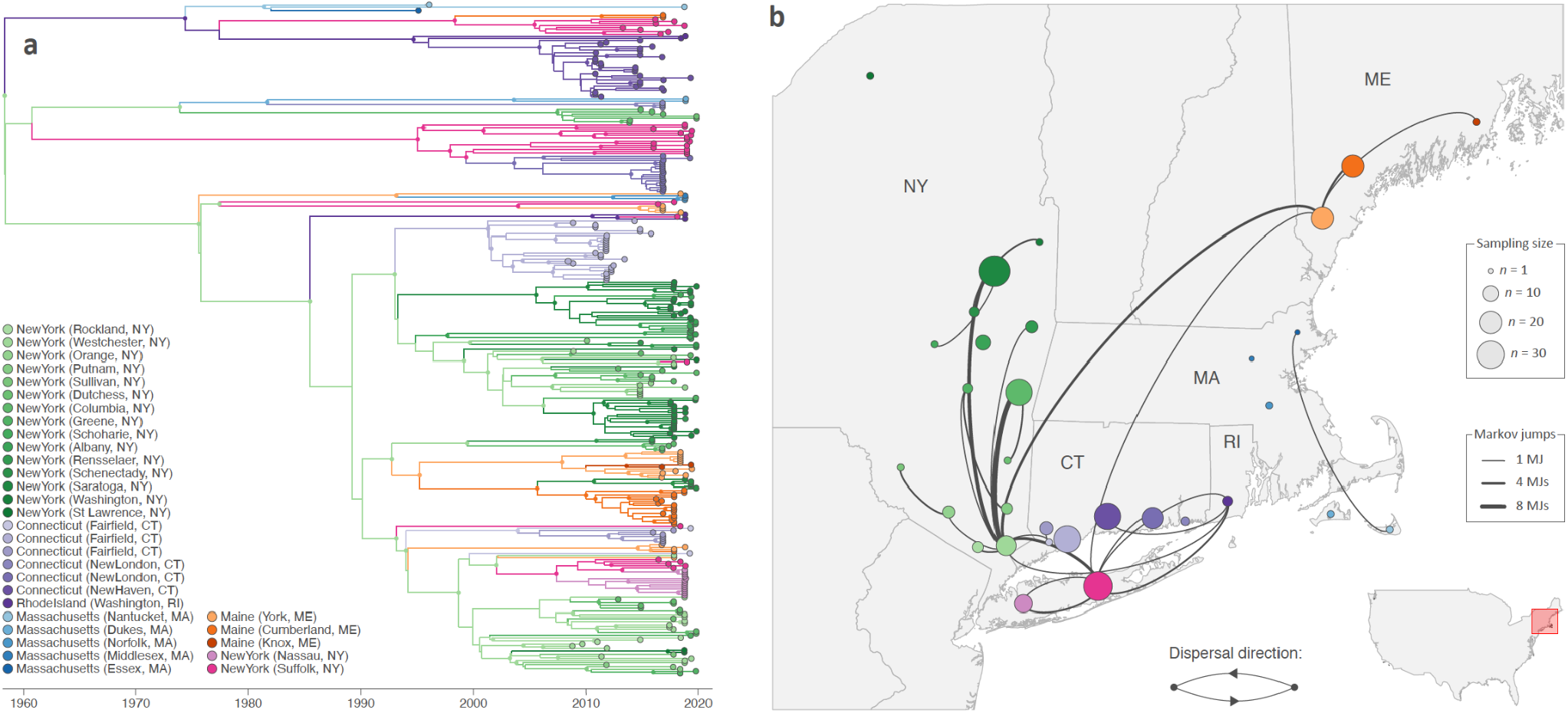
Discrete phylogeographic analysis of the dispersal history of Powassan virus in the northeastern United States. **(a)**Maximum clade credibility (MCC) tree with branches colored according to the locations inferred at the ancestral nodes. Tip nodes are colored according to their sampling location, and we only displayed internal nodes, using smaller dots and colored according to their inferred location, if they are associated with a posterior probability >0.95. **(b)**Sampling map and well-supported Markov jumps inferred by discrete phylogeographic inference. Sampling locations are displayed by dots with the size being proportional to the number of Powassan virus genomic sequences sampled and included in our analyses. We only report Markov jumps associated with an adjusted Bayes factor support higher than 3, which corresponds to positive support according to the scale of interpretation as previously defined (22).

### Distinct transmission foci

Like tick-borne encephalitis virus in Europe (30, 31), Powassan virus is hypothesized to be primarily maintained within strict foci (32, 33), meaning that the virus does not routinely migrate between locations. To test this hypothesis, we examined our discrete phylogeographic reconstruction of Powassan virus lineage II in the Northeast (**Fig. 2**). We found that sequences strongly cluster by location (**Fig. 2a**) with relatively few transition events over the past 20 years (**Fig. 2b**).

We further explored the spatial structure using a continuous phylogeographic approach (**Fig. 3**). These findings again highlight the highly focal distribution, with rare long-distance dispersal events leading to the establishment of new foci (**Fig. 3a-d**). These transmission foci are particularly clear in Connecticut, where, besides a single dispersal event between locations 1 (Westport) and 2 (Redding), we do not observe mixing between the 5 distinct locations (**Fig. 3d, inset**). For example, we estimate that the lineage II viruses sequenced in location 2 (Redding) have been separated from location 3 (Bridgeport) for ~33 years (95% HPD interval: 24-41), despite being less than 20 km apart.

**Fig. 3:**
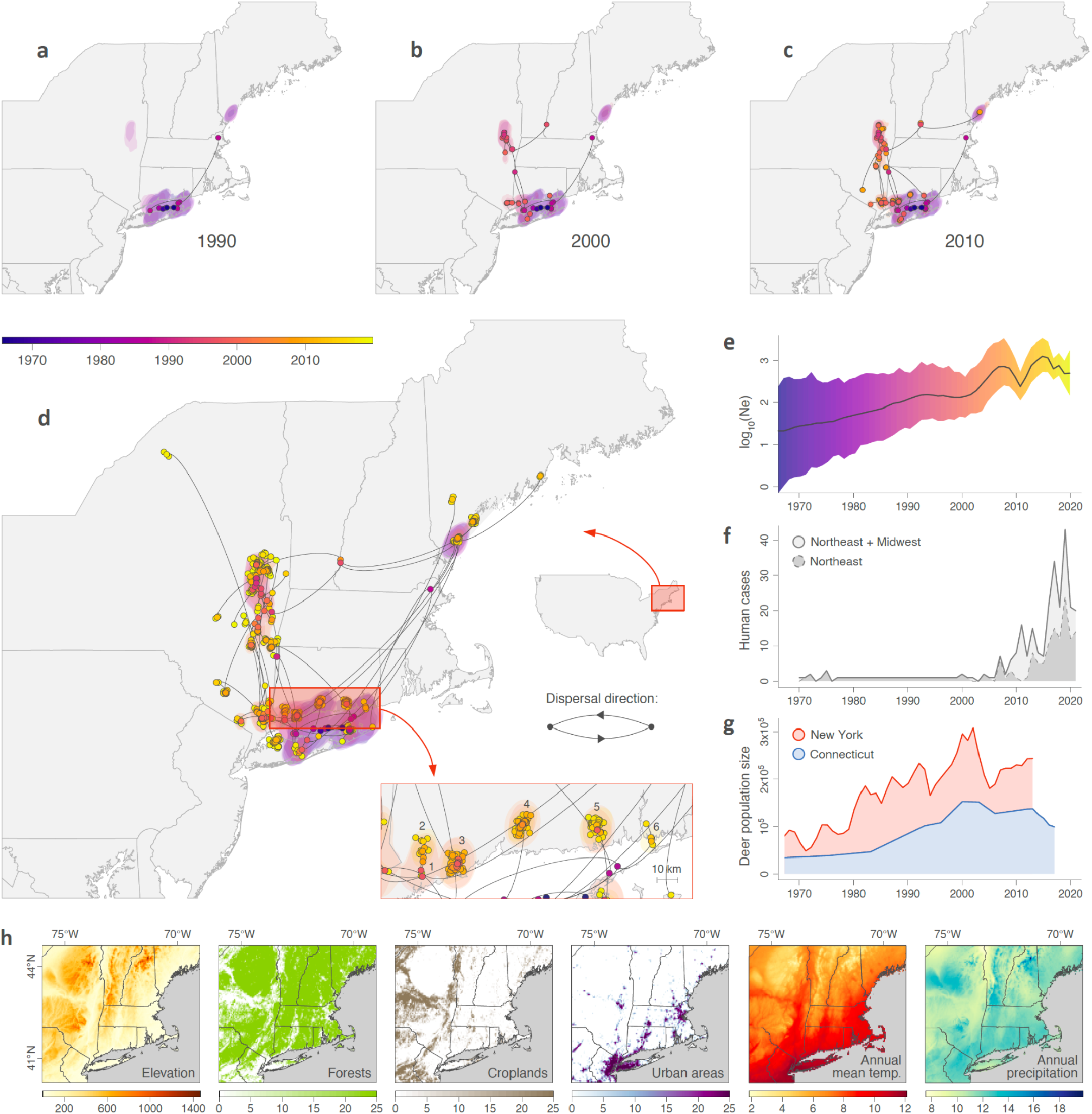
Spatially-explicit phylogeographic analysis of the dispersal history of Powassan virus in the northeastern United States. **(a-d)**Reconstruction of the dispersal history of Powassan virus lineages inferred by a spatially-explicit phylogeographic analysis. We mapped branches of the maximum clade credibility (MCC) tree reported in Figure 2 and whose nodes, as well as associated 80% highest posterior density (HPD) regions, are colored according to their time of occurrence. **(e)**skygrid reconstruction of the evolution of the overall effective size of the viral population (Ne). **(f)**Evolution of the number of confirmed human cases. **(g)**Estimation of the white-tailed deer (*Odocoileus virginianus*) populations in the states of New York and Connecticut. **(h)**Environmental factors included in landscape phylogeographic analyses to test their impact on Powassan virus dispersal.

We found the same pattern of isolated transmission foci with limited mixing for New York and Maine, albeit with more short distance dispersal events, particularly in New York (**Fig. 2b**, **Fig. 3d**). These differences may be explained due to differences in sampling (e.g. Powassan virus sequenced from ticks in New York included adults collected from deer), environmental barriers to spread (e.g. separation of Connecticut locations by rivers), or other ecological factors.

Overall, our data suggest that after the initial emergence of Powassan virus lineage II in the Northeast, migration between nearby and long-distance locations was relatively rare. This supports the hypothesis that Powassan virus is primarily maintained in highly localized transmission foci.

### Dispersal history

While we have shown that Powassan virus lineage II in the Northeast is primarily restricted to strict foci, we wanted to better understand the patterns and velocity of spread. Our spatially explicit continuous phylogeographic analysis indicates that Powassan virus lineage II emerged in the northeastern U.S. mostly following a south to north pattern (**Fig. 3a-d**). We estimate that the virus first became established in southern New York and Connecticut by the late 1950s (1940.3-1974.7; **Fig. 3a**). This was followed by a few long-distance dispersal events to more northern regions, perhaps by infected ticks feeding on birds which can migrate over longer distances. We estimate that the virus finally became established in Maine by 1991 (95% HPD = [1968-2011]) through multiple introductions. Our estimates of the relatively recent dispersion of Powassan virus lineage II in the northern part of the Northeast suggest that the virus is likely still emerging in parts of North America, following the northward expansion of *I. scapularis* (34).

We then used our continuous phylogeographic results to estimate the dispersal velocity of Powassan virus lineage II through the northeastern U.S. We estimated a weighted lineage dispersal velocity of ~3 km/year (95% HPD = [2.6-3.8]), which corresponds to a relatively slow dispersal capacity when compared to the estimates of the same metric obtained from the continuous phylogeographic analysis of other viruses (**Table S4**). For instance, Powassan lineage II dispersed faster than the rodent-borne Lassa virus in western Africa (~1 km/year) (35), but considerably slower than the mosquito-borne West Nile virus when it invaded North America (~165 km/year (36)).

### Population size

We next investigated if Powassan virus transmission has increased since its emergence in the Northeast, which could be a cause of the recent increase in reported human cases. We approached this by estimating the recent evolution of the virus effective population size through a ‘skygrid’ reconstruction, a non-parametric coalescent model assuming a panmictic population. Because the assumption of panmixia is never respected in practice, the effective population size (Ne) is sometimes interpreted as a measure of the overall genetic diversity of the viral population (e.g. (37)); although it has also been argued that such interpretation could be a bit reductive (38). Since the emergence in the Northeast, we found an overall increase in the effective population size of Powassan virus lineage II, but with growth stagnating since ~2005 (**Fig. 3e**). The latter could be a reflection of the virus becoming established across all of our study sites whereas the effective population size may still be increasing if we included additional sites in new emergence zones. Still, our data suggest that the increase in reported human cases in the region since 2010 (**Fig. 3f**) does not coincide with an increased virus effective population size (**Fig. 3e**). Thus, our findings do not support the hypothesis that the recent uptick in human cases is due to a significant increase in Powassan virus transmission; rather, it may be caused by an increase of human exposure to infected ticks.

As hypothesized as a significant factor for the timing of Powassan virus lineage II emergence in the Northeast, our overall estimates for the virus effective population size follow a similar trend as the population expansion of white-tailed deer in Connecticut and New York (**Fig. 3g**). Reforestation in the Northeast has led to dramatic increases in the white-tailed deer populations, followed by population expansion of *I. scapularis*, which in turn has facilitated the emergence of tick-borne pathogens such as *Borrelia burgdorferi* and *Babesia microti* (28). Our findings suggest that the cascading effect of population expansion of white-tailed deer and *I. scapularis* populations may also have facilitated the emergence of Powassan virus in the Northeast.

### Impact of environmental factors on the dispersal dynamic

We exploited our spatially-explicit phylogeographic reconstruction to investigate the impact of environmental factors on the dispersal dynamic of Powassan lineage II. As detailed in the Methods section, we tested the association between a series of environmental factors (**Fig. 3h**) and the dispersal location (39) as well as dispersal velocity of viral lineages (40). Our analyses revealed that inferred Powassan virus lineage II tended to avoid circulating in areas associated with relatively higher elevation (Bayes factor [BF] > 20; **Table S5**). However, outcomes of our analysis are strongly influenced by the sampling effort and pattern, and therefore we are able to describe environmental conditions related to dispersal locations of inferred viral lineages, but we cannot draw conclusions on the actual impact of those conditions on the dispersal (39). This observation could be related to higher abundance of *I. scapularis* at lower altitudes. Next, we investigated the impact of environmental factors on the dispersal velocity of Powassan virus lineage II. Our analyses did not highlight any environmental factor associated with the heterogeneity of Powassan lineage dispersal velocity across the study area (**Table S6**). This means that none of the tested factors increased the correlation between dispersal duration and geographic distance, the latter thus remaining the main resistance factor to dispersal.

### Discussion

Despite a rapid increase in the number of Powassan virus infections in humans over recent years, very little was known about the patterns of virus emergence and spread. By sequencing 279 Powassan virus genomes and using phylogeographic approaches, we have uncovered the patterns of virus emergence, transmission, and spread in the northeastern U.S. Our analyses revealed that Powassan virus lineage II likely emerged in the Northeast around 1940-1975, following the population growth of white-tailed deer and expansion of *I. scapularis* tick populations (10, 28). Powassan virus lineage II is maintained in highly localized transmission foci, with few migration events between relatively nearby locations. Our continuous phylogeographic analysis revealed that Powassan virus lineage II likely emerged from southern Connecticut into more northern regions with a weighted lineage dispersal velocity of ~3 km/year. Although we found an overall upward trend in the virus effective population size over the last decades, the recent increase in reported human cases of Powassan virus infection does not coincide with a higher effective population size. This suggests that the recent uptick in human cases is likely not due to a significant increase in Powassan virus enzootic transmission, but it may rather be due to other factors such as increased human exposure to infected ticks as well as an increase in case recognition. Our findings provide important insights in the local emergence patterns and transmission dynamics of Powassan virus in the northeastern U.S. Insights into the highly localized transmission foci that sustain Powassan virus transmission across multiple years, will help to identify areas with high risk of spillover to the human population, which can be targeted for prevention education or control efforts.

Our reconstruction of the emergence of Powassan virus in the northeast follows similar patterns as the emergence reported for other tick-borne pathogens such as *Borrelia*, *Babesia*, and *Anaplasma (41)*. Similar to Powassan virus, these pathogens are maintained in transmission cycles involving *I. scapularis* as the main vector and white-footed mice (*Peromyscus leucopus*) as the main host. Introduction of these tick-borne pathogens in the northeastern U.S. follows the reforestation in the 20th century leading to rapid population expansions of both white-tailed deer and *I. scapularis (28, 34)*. Despite the similarities in emergence and ecology, other tick-borne pathogens seem to have more widespread distributions as compared to the highly focal distribution of Powassan virus. Previous studies on *Borrelia burgdorferi* reported high genetic diversity within local populations (42), lack of population genetic structure (25), and no strict genetic clustering by location within the Northeast (11). This suggests that despite the similarities in ecology, *B. burgdorferi* and Powassan virus are maintained by different mechanisms.

To explain the highly focal distribution, we have formulated two main hypotheses on how Powassan virus lineage II may be maintained in strict transmission foci (**Fig. 4**). Our first hypothesis is that vertical transmission from adult to the next larval stage plays a minor role in the Powassan virus transmission cycle. Adult *I. scapularis* ticks preferentially feed on larger mammals, such as white-tailed deer, and we would expect more mixing of Powassan virus clades if infected adults travel across larger distances when feeding on deer, particularly during early years of emergence when deer habitats were less fragmented. Inefficient vertical transmission from adult to larvae would explain why adult ticks and deer may move, while Powassan virus foci remain local. Moreover, low Powassan virus infection rates in ticks further reduce the probability of successful establishment in new areas (43). Previous studies have provided evidence for vertical transmission of Powassan virus (i.e. one of six infected females transmitted Powassan virus to its progeny) (44), but it remains unclear what percentage of progeny within an egg batch becomes infected. Future studies can help to test this hypothesis by determining rates of vertical transmission in the laboratory, and determining infection rates of unfed larvae in the field.

**Fig. 4:**
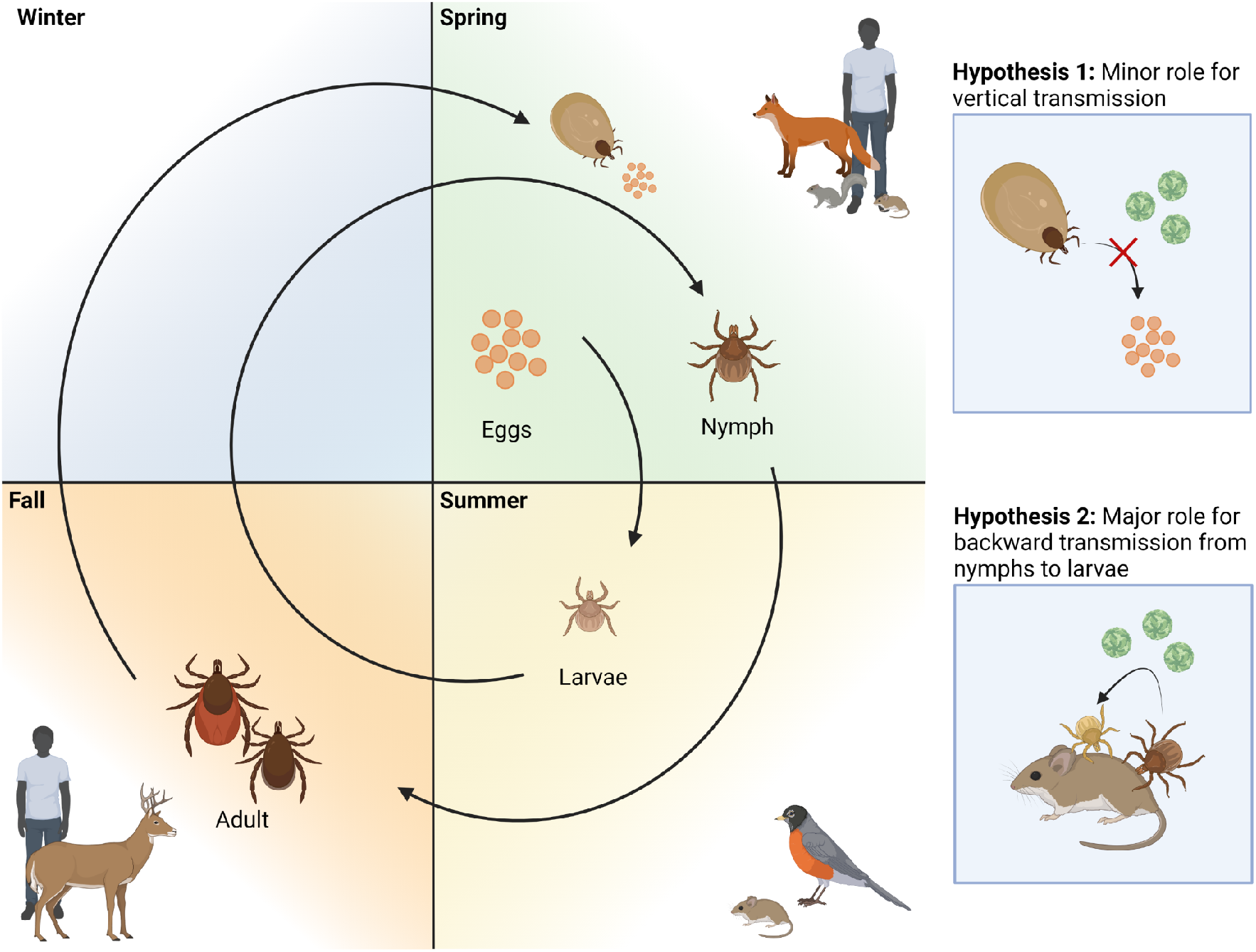
*Ixodes scapularis* life cycle with proposed hypotheses that may explain the focal distribution of Powassan virus. Life cycle adapted from (41) and created with bioRender.com. Two hypotheses are proposed that may explain the focal distribution of Powassan virus.

Our second hypothesis is that backward transmission through tick life stages (e.g. from nymphs to larvae or adults to nymphs) is the main mechanism for Powassan virus maintenance (**Fig. 4**). Powassan virus clades better match the strong geographical structure of white-footed mice (45, 46) and other small mammals that share the same habitat rather than the weak structure of *I. scapularis* ticks (26). This suggests that Powassan virus is mostly maintained by local transmission cycles that include larvae and nymphs feeding on small mammals, and adult ticks and other animals that can travel long distances are primarily dead-ends, although sporadic long-distance dispersal events may happen. Backward transmission from nymphs to larvae may either occur through direct co-feeding transmission in the absence of systemic infection of the rodent host, as has been reported for tick-borne encephalitis virus (47, 48), or through feeding on viremic hosts. To further investigate this hypothesis, we propose studies to determine the frequency of co-feeding between nymphs and larvae, to determine the size of transmission foci, and to perform comparative genetics between Powassan virus, *I. scapularis*, and small mammalian hosts, such as white-footed mice. Additionally, further research is needed to better understand the drivers and frequency of long distance dispersal events that may result in the establishment of new transmission foci. Future ecological and genetic studies on host (e.g. bird and deer) and tick population movement are needed to better understand how movement of both host and vector may contribute to the occasional long distance dissemination of Powassan virus.

Our study has several limitations. First, sampling strategy was not standardized across the Northeastern United States nor was it standardized across time. Powassan virus surveillance activities are organized at the state level, and therefore there are differences in effort between states. For instance, samples sequenced from Connecticut and Maine were collected from the same sites during multiple years, whereas a much larger number of varying sites were sampled from year to year in New York. Sampling in New York also included tick collections from hunter-harvested deer, whereas sampling in Connecticut and Maine was exclusively done via drag sampling. Spatial heterogeneity in sampling effort will impact phylogeographic reconstructions, with unsampled locations being excluded from the reconstructed dispersal history, and undersampling of locations may lead to underestimation of the degree of connectivity between sites. Second, we attempted to include as many publicly available genomes in our analyses, but some genomes had to be excluded because they did not have complete metadata (e.g. missing collection date and location) or because they did not fit the molecular clock (e.g. Powassan sequences from human infections). Lastly, choice of molecular clock model may impact estimates for the evolutionary rate and the tMRCA. For example, when performing the analyses with a strict molecular clock model instead of a relaxed clock model, we get a lower estimated evolutionary rate of 3.76×10^−5^ substitutions/site/year (95% HPD = 2.59-5.45×10^−5^) and an earlier tMRCA of 1881.3 (95% HPD = 1832.4-1934.8). This highlights the importance of molecular clock model choice based on best model fit and that we should be careful in our interpretation of the relaxed clock model estimates. Our estimates of emergence before the mid 1970s should be interpreted as emergence latest by this time period, and not as an exact estimate of the emergence time.

Our study has important implications for our understanding of Powassan virus transmission dynamics and future control. Currently, no vaccines or specific treatments are available for Powassan virus infection, which leaves prevention of disease highly dependent on education and control. Our estimates of the dispersal velocity can help to better inform the risk of regions in proximity of the current distribution of Powassan virus in the Northeast, which may be targeted for tick and virus surveillance as well as potential future control. Furthermore, the identification of highly localized transmission foci provides both opportunities for better education of the general public about high risk areas and effective targeted control in Powassan virus hotspots. Eradication of Powassan virus in transmission foci that have been maintained for several years without introductions of new virus clades, will likely result in highly effective and long lasting control.

## Methods

### Sample collection

Tick collections, nucleic acid extraction, and Powassan virus screening were done at the Connecticut Agricultural Experiment Station (CAES), MaineHealth Institute for Research (MHIR), New York State Department of Health (NYSDOH), and Cornell University (32, 43, 49–51). Briefly, the majority of ticks were collected by dragging a white cloth over the ground and low vegetation, and a smaller proportion were collected from vertebrate hosts (e.g. deer). All collected ticks were sorted by species, life stage, collection site and date before screening for pathogens. Individual or pooled ticks were homogenized and nucleic acid was extracted according to manufacturer’s instructions (**Table S1**). Powassan virus was detected by RT-qPCR or nanochip assays (32, 43, 49–51). Samples collected from 2008 through 2016 in Connecticut were passaged once on BHK-21 cells before nucleic acid extraction.

### Untargeted metagenomic Illumina sequencing

We initially used an untargeted metagenomic approach to sequence Powassan virus isolates which were passaged on cells at CAES. Our protocol is adapted from (52), and is openly available (53). In brief, 10 μL of extracted nucleic acid was treated with DNase I (New England Biolabs, Ipswich, MA), followed by a clean up step using a ratio of 1.8:1 beads to sample. All clean up steps were done using MagBind TotalPure NGS magnetic beads (Omega Biotek, Norcross, GA) with automated protocols for the Kingfisher flex purification system (Thermo Fisher Scientific, Waltham, MA). First-strand cDNA was synthesized using SuperScript IV VILO (Thermo Fisher Scientific) and second-strand cDNA using *Escherichia coli* DNA ligase and polymerase (New England Biolabs), followed by a clean up step (1.8:1 beads to sample ratio). Libraries were prepared using the Nextera XT DNA library preparation kit for Illumina (Illumina, San Diego, CA), according to manufacturer’s instructions but using less than the recommended reagent volumes (54). Individual and pooled libraries were quantified using the 1x dsDNA HS assay kit on the Qubit 4 (Thermo Fisher Scientific) and size distribution was determined using the high sensitivity DNA kit on the Bioanalyzer 2100 (Agilent, Santa Clara, CA). Pooled libraries were sequenced on the Illumina NovaSeq (paired-end 150) at the Yale Center for Genome Analysis. PCR duplicates were removed, reads were aligned to the reference genome using Bowtie2, and consensus genomes were called at a minimum frequency threshold of 0.75 and minimum coverage of 10X using Geneious Prime 2020.0.4.

### Targeted amplicon-based sequencing

Although we were able to successfully sequence Powassan virus from cell culture-passage samples using untargeted metagenomics, we developed an amplicon-based sequencing approach to improve coverage when sequencing from tick homogenates. We used an adapted protocol (55), using Nextera XT for library prep as developed for SARS-CoV-2 amplicon-based sequencing (56). In brief, cDNA was synthesized from 10 μL of RNA using SuperScript IV VILO (Thermo Fisher Scientific). Two separate primer pools were prepared by mixing equal volumes of each primer with a concentration of 10 μM (**Table S2**). The two primer pools were used to generate tiled amplicons using Q5 high-fidelity 2X master mix (New England Biolabs), followed by a clean up step and quantification using the Qubit. Amplicons were diluted to 1 ng/μL and combined for library prep as described above. Pooled libraries were quantified on Qubit and size distribution were determined on the bioanalyzer. Pooled libraries were sequenced on the Illumina NovaSeq (paired-end 150) at the Yale Center for Genome Analysis. Consensus genomes were generated at a minimum frequency threshold of 0.75 and minimum coverage of 10X using iVar version 1.2.3. All sequencing data is publicly available under BioProject PRJNA889421 and **Table S3**.

### Powassan virus phylogeny

We sequenced 279 Powassan virus genomes and estimated a maximum-likelihood tree using IQ-TREE version 1.6.12 with ultrafast bootstrap approximation (1,000 replicates) (57) to determine phylogenetic relationships between publicly available and newly sequenced Lineage I and II genomes.

### Temporal signal assessment

To evaluate if our Powassan virus data set contains sufficient temporal signal and would permit time-calibrated analyses using molecular clock models (and hence constitutes a measurably evolving population), we performed a Bayesian Evaluation of Temporal Signal (BETS) analysis (21, 58). This analysis involves assessing the model fit to the data of both a strict clock and an uncorrelated relaxed clock with an underlying lognormal distribution, both with and without the sampling dates associated with the genomes in our data set (**Table 1**). We employed generalized stepping-stone sampling (59) to accurately estimate the log marginal likelihood of each of these four models. For each log marginal likelihood estimation, we ran an initial Markov chain of 500 million iterations, followed by 500 power posteriors that are explored during 1 million iterations, logging every 1000 iterations. We used the following priors: a lognormal prior distribution (mean = 1.0, standard deviation = 1.25) on the transition-transversion ratio ***κ***; a Dirichlet prior distribution (1.0; K = 4) on the nucleotide base frequencies; an exponential prior distribution (mean = 0.5) on the shape parameter describing the discretized gamma distribution used to model among-site rate heterogeneity; a gamma prior distribution (shape = 0.001; scale = 1000.0) on the constant population size; a CTMC reference prior (60) on the (constant) evolutionary rate under the strict clock model; a CTMC reference prior (60) on the mean evolutionary rate of the underlying lognormal distribution for the uncorrelated relaxed clock model; an exponential prior distribution (mean = 1/3) on the standard deviation of the underlying lognormal distribution for the uncorrelated relaxed clock model. BEAST “.xml” files for the BETS analysis are available on github (https://github.com/grubaughlab/powassan-genomics/).

### Discrete phylogeographic reconstruction

To investigate the dispersal history of Powassan virus lineages in the northeastern U.S., we first conducted a discrete phylogeographic analysis using the Bayesian stochastic search variable selection (BSSVS) model (61) implemented in BEAST 1.10 (62). For this analysis, we considered each U.S. county of origin as a distinct location, except for the Connecticut area where each sampling site was considered as a distinct discrete location. We modeled the branch-specific evolutionary rates according to a relaxed molecular clock with an underlying log-normal distribution (63) and the nucleotide substitution process according to a GTR+Γ parameterisation (64); and we specified a skygrid coalescent model as the tree prior (65). Both the substitution and coalescent models were selected for their flexibility. Three independent Markov chain Monte Carlo (MCMC) algorithms were run for 5×10^8^ iterations and sampled every 10^5^ iterations. Resulting posterior distributions were eventually combined after having discarded 10% of sampled trees in each of them. We used the program Tracer 1.7 (66) for assessing the convergence and mixing properties, and that estimated sampling size (ESS) values associated with estimated parameters were all >200 after having combined the outputs of the three independent analyses. We then used the program TreeAnnotator 1.10 (62) to obtain a maximum clade credibility (MCC) tree. We reported Markov jumps between discrete locations as estimated by the BSSVS analyses and supported by an adjusted Bayes factor (BF) values >3, which correspond to at least “positive” statistical support following the scale of interpretation defined by Kass & Raftery (22). The adjusted BF support takes into account the relative abundance of samples by location (67) and is based on a tip labels swapping procedure similar to a tip date randomization that can be performed to test for temporal signal (68). BEAST “.xml” files and concatenated log files for the discrete phylogeographic analysis are available on github (https://github.com/grubaughlab/powassan-genomics/).

### Continuous phylogeographic reconstruction

To reconstruct the dispersal history of Powassan virus lineages in a spatially-explicit context, we performed a continuous phylogeographic analysis using the relaxed random walk (RRW) diffusion model (69, 70) implemented in the software package BEAST 1.10 (62), with a gamma distribution to model the among-branch heterogeneity in diffusion velocity. As for the discrete phylogeographic analysis, branch-specific evolutionary rates were modeled according to a relaxed molecular clock with an underlying log-normal distribution and the nucleotide substitution process according to a GTR+Γ parameterisation (64); and we also specified a flexible skygrid model as the tree prior (65). The Markov chain Monte-Carlo (MCMC) algorithm was run for 15×10^8^ generations and sampled every 10^5^ generations. We used the program Tracer 1.7 for assessing the convergence and mixing properties, and that estimated sampling size (ESS) values associated with estimated parameters were all >200, the program TreeAnnotator 1.10 to obtain a maximum clade credibility (MCC) tree, as well as the R package “seraphim” (71, 72) to extract the spatiotemporal information embedded within posterior trees and to estimate the weighted lineage dispersal velocity, the latter being defined as follows:

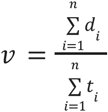

where *d_i_* and *t_i_* are the geographic distance traveled by the phylogeny branch and the duration of that branch, respectively. BEAST “.xml” files and concatenated log files for the continuous phylogeographic analysis are available on github (https://github.com/grubaughlab/powassan-genomics/).

### Landscape phylogeographic analyses

Landscape phylogeographic analyses aim at exploiting phylogeographic reconstructions to unravel the impact of environmental factors on the dispersal history and dynamic of viral lineages (73). Specifically, we implemented two previously introduced analytical procedures to investigate the impact of environmental factors on the dispersal location (39) and velocity (40) of Powassan virus lineages. Both analytical procedures here rely on the comparison between inferred and randomized spatially-annotated trees, the latter sharing the time-scaled topology of the inferred trees but with phylogenetic branch positions that had been randomized across the study area. In practice, phylogenetic node positions were randomized within the study area, under the constraint that branch length (i.e. geographic distance connecting both branch nodes), branch duration, tree topology, and root position remained unchanged (71). The purpose of these randomizations is thus to obtain spatially-annotated trees corresponding to the trees inferred by continuous phylogeography but along which we generated a new diffusion process that has been impacted by any environmental factor.

We first investigated whether Powassan virus lineages tended to avoid or preferentially circulate within areas associated with particular environmental conditions. For this purpose, we extracted and subsequently averaged the environmental values at the tree node positions to obtain, for each environmental factor, a posterior distribution of mean environmental values at tree node positions. We then compared values obtained through inferred trees and their corresponding randomized trees by approximating a BF with the following formula (74): BF = (*p_e_*/(1-*p_e_*))/(0.5/(1-0.5)). To test if a particular environmental factor *e* tended to attract viral lineages, *p_e_* was defined as the frequency at which the environmental values from inferred trees were greater than values from randomized trees; and to test if a particular environmental factor *e* tended to repulse viral lineages, *p_e_* was defined as the frequency at which the environmental values from inferred trees were lower than values from randomized trees. We considered BF values > 20 and 3 < BF < 20 as strong and positive statistical supports, respectively (22).

Second, we investigated to what extent Powassan virus lineage dispersal velocity was impacted by environmental factors acting as conductance or resistance factors. For each branch in the inferred and randomized trees we calculated an “environmental distance” using two path models: the least-cost path (75) and Circuitscape (76) algorithms, the latter accommodating uncertainty in the travel route. An environmental distance is calculated first from the raster of the environmental variable, and second from a uniform “null” raster whose cell values are all set to “1”. The environmental distance is a spatial distance that is weighted according to the values of the underlying environmental raster, and therefore constitutes a proxy for geographical distance when computed on the null raster. Each environmental variable was considered twice: once as a potential conductance factor that facilitates movement, and once as a potential resistance factor that impedes it. For each environmental variable, we also generated and tested several distinct rasters by transforming the original raster cell values with the following formula: *v_t_* = 1 + *k*(*v_o_*/*v_max_*), where *v_t_* and *v_o_* are the transformed and original cell values, and *v_max_* the maximum cell value recorded in the raster. The rescaling parameter *k* here allows the definition and testing of different strengths of raster cell conductance or resistance, relative to the conductance/resistance of a cell with a minimum value set to “1”, which corresponds to the “null” raster. For each environmental variable, we generated three distinct rasters using the following values for rescaling factor *k*: *k* = 10, 100, and 1000. For these analyses, we estimated the statistic *Q* defined as the difference between the coefficient of determinations obtained (i) when branch durations are regressed against the environmental distances computed on an environmental and (ii) when branch durations are regressed against the environmental distances computed on the null raster. We estimated a *Q* statistic for each environmental raster and each of the 1,000 trees sampled from the posterior distribution. An environmental factor was only considered as potentially explanatory if both its distribution of regression coefficients and its associated distribution of *Q* values were positive (77), i.e. with at least 90% of positive values. In this case, the statistical support associated with the resulting *Q* distribution was compared with the corresponding null of distribution of *Q* values obtained when computing environmental distances for phylogenetic branches of randomized trees. Similar to the procedure used for the investigation of the impact of environmental factors on the dispersal locations of Powassan virus lineages, the comparisons between inferred and randomized distributions of *Q* values was formalized by approximating a Bayes factor support (40).

## Data availability

All data are included in this article, the SI Appendix, github (https://github.com/grubaughlab/powassan-genomics/), and in BioProject PRJNA889421.

## Acknowledgements

We would like to thank Anne Piantadosi, Erica Normandin, Pardis C. Sabeti, Rebekah McMinn, Greg Ebel, Heidi Goethert, Sam R. Telford III, Sebastian Lequime, Alexander A. Fisher, and Marc A. Suchard for their input in discussions on the methods and results of this study.

## Funding

This publication was made possible by CTSA Grant Number UL1 TR001863 from the National Center for Advancing Translational Science (NCATS), a component of the National Institutes of Health (NIH) awarded to CBFV, and the National Institute Of Allergy And Infectious Diseases of the NIH under Award Number 1R56 AI149004-01A1 awarded to NDG. Its contents are solely the responsibility of the authors and do not necessarily represent the official views of NIH. GB acknowledges support from the Internal Funds KU Leuven under grant agreement C14/18/094 and the Research Foundation - Flanders (“Fonds voor Wetenschappelijk Onderzoek - Vlaanderen,” G0E1420N, G098321N).

## Author contributions

CBFV, SD, NDG designed the study; DEB, APD, RMR, SCW, JFA, CBL, REL, MAP, LDK, JLG-K, LBG, RPS, PMA, and ATC collected data/samples; CBFV and JRF performed sequencing; CBFV, AFB, GB, SD, NDG analyzed the data; CBFV, SD, and NDG drafted the manuscript; all authors reviewed and approved the manuscript.

## Competing interests

NDG is a consultant for Tempus Labs and the National Basketball Association for work related to COVID-19. All other authors declare no competing interests.

---

**Table S1:**
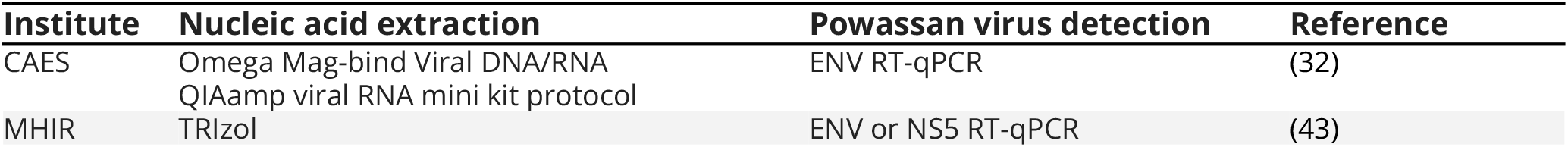
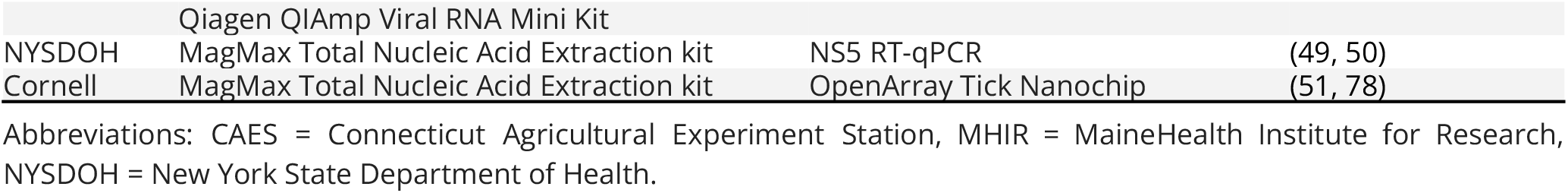
Nucleic acid extraction and Powassan screening methods used by institutes providing samples.

**Table S2:**
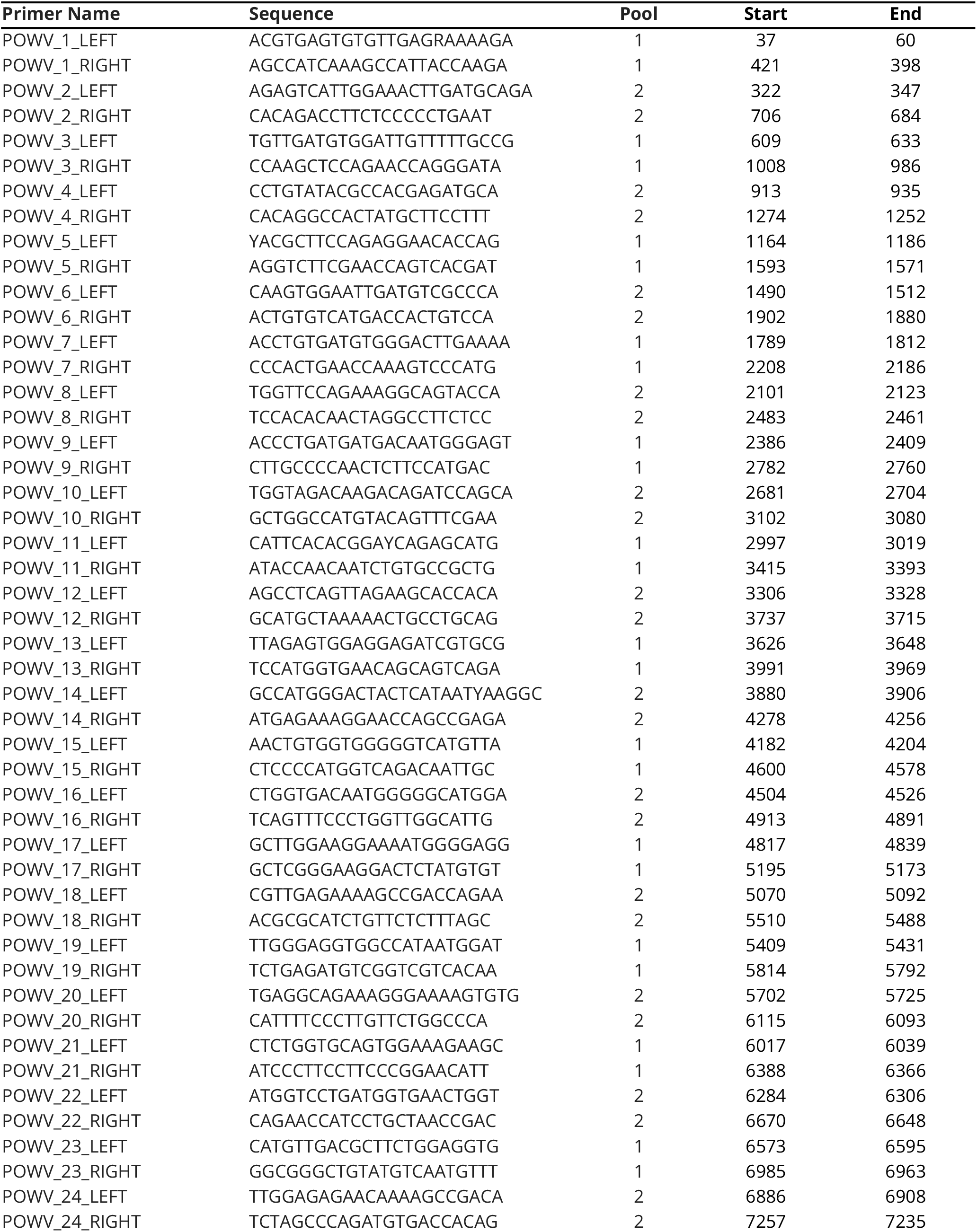
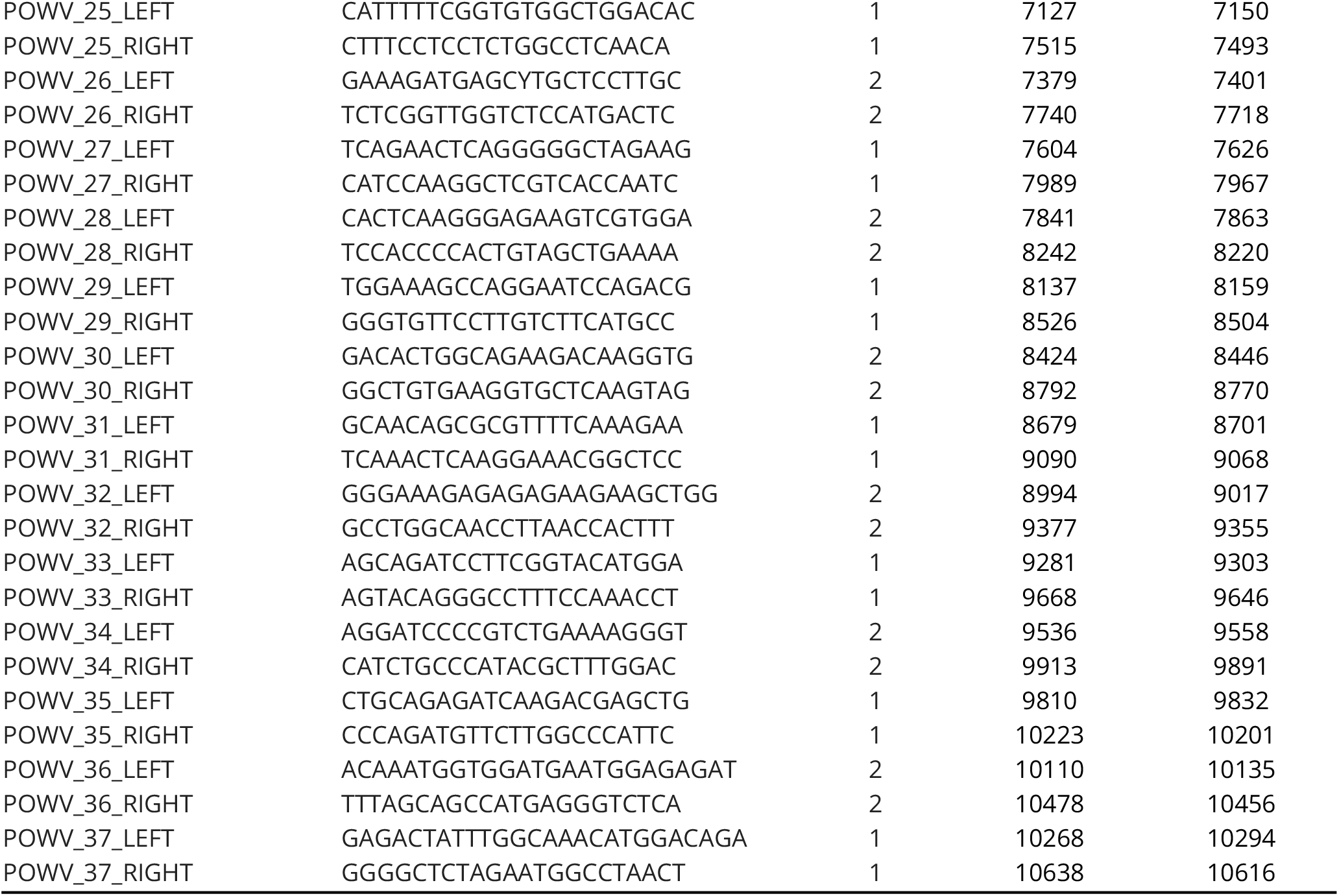
PrimalSeq primer scheme for Powassan virus amplicon-based sequencing.

**Table S3:** Newly sequenced Powassan virus genomes.

| Source | Sample | Accession |
| --- | --- | --- |
| CAES | PV001-P1734_LII_Bridgeport_CT_USA_Ixodes-scapularis_F_Drag_2014-10-29 | OL704116 |
| CAES | PV002-P1735_LII_Bridgeport_CT_USA_Ixodes-scapularis_M_Drag_2014-10-29 | OL704117 |
| CAES | PV003-P1663_LII_North-Branford_CT_USA_Ixodes-scapularis_F_Drag_2014-10-30 | OL704118 |
| CAES | PV004-P1665_LII_North-Branford_CT_USA_Ixodes-scapularis_F_Drag_2014-10-30 | OL704119 |
| CAES | PV005-P1666_LII_North-Branford_CT_USA_Ixodes-scapularis_M_Drag_2014-10-30 | OL704120 |
| CAES | PV006-P1677_LII_North-Branford_CT_USA_Ixodes-scapularis_F_Drag_2014-10-30 | OL704121 |
| CAES | PV007-P1384_LII_Bridgeport_CT_USA_Ixodes-scapularis_F_Drag_2014-05-02 | OL704122 |
| CAES | PV008-P1429_LII_North-Branford_CT_USA_Ixodes-scapularis_N_Drag_2014-06-24 | OL704123 |
| CAES | PV009-P1742_LII_North-Branford_CT_USA_Ixodes-scapularis_F_Drag_2014-10-27 | OL704124 |
| CAES | PV010-P1783A_LII_North-Branford_CT_USA_Ixodes-scapularis_F_Drag_2014-10-30 | OL704125 |
| CAES | PV011-P1774A_LII_North-Branford_CT_USA_Ixodes-scapularis_F_Drag_2014-10-28 | OL704126 |
| CAES | PV012-P913_LII_Bridgeport_CT_USA_Ixodes-scapularis_N_Drag_2013-06-12 | OL704127 |
| CAES | PV013-P2552_LII_Bridgeport_CT_USA_Ixodes-scapularis_M_Drag_2015-10-22 | OL704128 |
| CAES | PV014-8_LII_Redding_CT_USA_Ixodes-scapularis_F_Drag_2015-12-28 | OL704129 |
| CAES | PV015-67_LII_North-Branford_CT_USA_Ixodes-scapularis_F_Drag_2015-12-03 | OL704130 |
| CAES | PV016-32_LII_Redding_CT_USA_Ixodes-scapularis_F_Drag_2016-11-10 | OL704131 |
| CAES | PV017-45_LII_Lyme_CT_USA_Ixodes-scapularis_F_Drag_2016-11-28 | OL704132 |
| CAES | PV018-50_LII_Lyme_CT_USA_Ixodes-scapularis_M_Drag_2016-11-28 | OL704133 |
| CAES | PV019-52_LII_Lyme_CT_USA_Ixodes-scapularis_M_Drag_2016-11-28 | OL704134 |
| CAES | PV020-53_LII_Lyme_CT_USA_Ixodes-scapularis_M_Drag_2016-11-28 | OL704135 |
| CAES | PV021-55_LII_Lyme_CT_USA_Ixodes-scapularis_F_Drag_2016-11-28 | OL704136 |
| CAES | PV022-62_LII_Lyme_CT_USA_Ixodes-scapularis_M_Drag_2016-11-16 | OL704137 |
| CAES | PV023-93_LII_Lyme_CT_USA_Ixodes-scapularis_F_Drag_2016-11-07 | OL704138 |
| CAES | PV024-95_LII_Lyme_CT_USA_Ixodes-scapularis_F_Drag_2016-11-07 | OL704139 |
| CAES | PV025-98_LII_Lyme_CT_USA_Ixodes-scapularis_F_Drag_2016-11-07 | OL704140 |
| CAES | PV026-101_LII_Lyme_CT_USA_Ixodes-scapularis_F_Drag_2016-11-07 | OL704141 |
| CAES | PV027-114_LII_Lyme_CT_USA_Ixodes-scapularis_M_Drag_2016-11-01 | OL704142 |
| CAES | PV028-116_LII_Lyme_CT_USA_Ixodes-scapularis_M_Drag_2016-11-01 | OL704143 |
| CAES | PV029-118_LII_Lyme_CT_USA_Ixodes-scapularis_M_Drag_2016-11-01 | OL704144 |
| CAES | PV030-122_LII_Lyme_CT_USA_Ixodes-scapularis_F_Drag_2016-11-01 | OL704145 |
| CAES | PV031-124_LII_Lyme_CT_USA_Ixodes-scapularis_F_Drag_2016-11-01 | OL704146 |
| CAES | PV032-125_LII_Lyme_CT_USA_Ixodes-scapularis_F_Drag_2016-11-01 | OL704147 |
| CAES | PV033-126_LII_Lyme_CT_USA_Ixodes-scapularis_F_Drag_2016-11-01 | OL704148 |
| CAES | PV034-P0375_LII_Bridgeport_CT_USA_Ixodes-scapularis_M_Drag_2010-11-01 | OL704149 |
| CAES | PV035-P0183_LII_Bridgeport_CT_USA_Ixodes-scapularis_F_Drag_2008-11-01 | OL704150 |
| CAES | PV036-P0096_LII_Bridgeport_CT_USA_Ixodes-scapularis_F_Drag_2008-11-01 | OL704151 |
| CAES | PV037-P0494_LII_Bridgeport_CT_USA_Ixodes-scapularis_F_Drag_2010-11-02 | OL704152 |
| CAES | PV038-P0525_LII_North-Branford_CT_USA_Ixodes-scapularis_F_Drag_2010-10-29 | OL704153 |
| CAES | PV039-P0526_LII_North-Branford_CT_USA_Ixodes-scapularis_F_Drag_2010-10-29 | OL704154 |
| CAES | PV040-P0618_LII_North-Branford_CT_USA_Ixodes-scapularis_F_Drag_2010-10-29 | OL704155 |
| CAES | PV041-P0621_LII_North-Branford_CT_USA_Ixodes-scapularis_F_Drag_2010-10-29 | OL704156 |
| CAES | PV042-P0629_LII_North-Branford_CT_USA_Ixodes-scapularis_M_Drag_2010-10-29 | OL704157 |
| CAES | PV043-P1697_LII_Bridgeport_CT_USA_Ixodes-scapularis_F_Drag_2011-11-09 | OL704158 |
| CAES | PV045-P0690_LII_North-Branford_CT_USA_Ixodes-scapularis_M_Drag_2011-04-26 | OL704159 |
| CAES | PV046-P0681_LII_North-Branford_CT_USA_Ixodes-scapularis_M_Drag_2011-04-26 | OL704160 |
| CAES | PV047-P0785_LII_North-Branford_CT_USA_Ixodes-scapularis_M_Drag_2011-05-09 | OL704161 |
| CAES | PV048-P0751_LII_Bridgeport_CT_USA_Ixodes-scapularis_F_Drag_2011-04-27 | OL704162 |
| CAES | PV049-P0760_LII_Bridgeport_CT_USA_Ixodes-scapularis_M_Drag_2011-04-27 | OL704163 |
| CAES | PV050-P0795_LII_North-Branford_CT_USA_Ixodes-scapularis_F_Drag_2011-05-09 | OL704164 |
| CAES | PV051-P0933_LII_North-Branford_CT_USA_Ixodes-scapularis_F_Drag_2011-10-21 | OL704165 |
| CAES | PV052-P1113_LII_Bridgeport_CT_USA_Ixodes-scapularis_M_Drag_2011-10-18 | OL704166 |
| CAES | PV053-P1104_LII_Bridgeport_CT_USA_Ixodes-scapularis_M_Drag_2011-10-18 | OL704167 |
| CAES | PV054-P1109_LII_Bridgeport_CT_USA_Ixodes-scapularis_F_Drag_2011-10-18 | OL704168 |
| CAES | PV055-P1112_LII_Bridgeport_CT_USA_Ixodes-scapularis_F_Drag_2011-10-18 | OL704169 |
| CAES | PV056-P1114_LII_Bridgeport_CT_USA_Ixodes-scapularis_M_Drag_2011-10-18 | OL704170 |
| CAES | PV057-P1107_LII_Bridgeport_CT_USA_Ixodes-scapularis_F_Drag_2011-10-18 | OL704171 |
| CAES | PV058-P1241_LII_Bridgeport_CT_USA_Ixodes-scapularis_F_Drag_2011-10-28 | OL704172 |
| CAES | PV059-P1514_LII_Bridgeport_CT_USA_Ixodes-scapularis_M_Drag_2011-10-25 | OL704173 |
| CAES | PV060-P1518_LII_Bridgeport_CT_USA_Ixodes-scapularis_F_Drag_2011-10-25 | OL704174 |
| CAES | PV061-P1730_LII_Bridgeport_CT_USA_Ixodes-scapularis_F_Drag_2011-11-09 | OL704175 |
| CAES | PV062-P1753_LII_Bridgeport_CT_USA_Ixodes-scapularis_M_Drag_2011-11-09 | OL704176 |
| CAES | PV063-P1912_LII_Bridgeport_CT_USA_Ixodes-scapularis_M_Drag_2012-04-05 | OL704177 |
| CAES | PV064-P1933_LII_Bridgeport_CT_USA_Ixodes-scapularis_F_Drag_2012-04-05 | OL704178 |
| CAES | PV065-P1930_LII_Bridgeport_CT_USA_Ixodes-scapularis_F_Drag_2012-04-05 | OL704179 |
| CAES | PV066-P1184_LII_North-Branford_CT_USA_Ixodes-scapularis_N_Drag_2014-05-29 | OL704180 |
| CAES | PV067-P1290_LII_North-Branford_CT_USA_Ixodes-scapularis_Mix_Drag_2014-05-29 | OL704181 |
| CAES | PV068-P1295_LII_North-Branford_CT_USA_Ixodes-scapularis_Mix_Drag_2014-05-14 | OL704182 |
| CAES | PV069-165_LII_Redding_CT_USA_Ixodes-scapularis_M_Drag_2016-11-10 | OL704183 |
| CAES | PV070-188_LII_North-Branford_CT_USA_Ixodes-scapularis_F_Drag_2016-12-06 | OL704184 |
| CAES | PV071-219_LII_Redding_CT_USA_Ixodes-scapularis_M_Drag_2016-11-14 | OL704185 |
| CAES | PV072-228_LII_Redding_CT_USA_Ixodes-scapularis_M_Drag_2016-11-14 | OL704186 |
| CAES | PV073-237_LII_Redding_CT_USA_Ixodes-scapularis_F_Drag_2016-11-14 | OL704187 |
| CAES | PV074-3338_LII_Groton_CT_USA_Ixodes-scapularis_F_Drag_2016-11-02 | OL704188 |
| CAES | PV075-3381_LII_Groton_CT_USA_Ixodes-scapularis_F_Drag_2016-11-02 | OL704189 |
| CAES | PV076-3464_LII_Groton_CT_USA_Ixodes-scapularis_F_Drag_2016-11-02 | OL704190 |
| CAES | PV077-3569_LII_North-Branford_CT_USA_Ixodes-scapularis_F_Drag_2016-10-17 | OL704191 |
| CAES | PV078-3591_LII_North-Branford_CT_USA_Ixodes-scapularis_M_Drag_2016-10-20 | OL704192 |
| CAES | PV080-160_LII_North-Branford_CT_USA_Ixodes-scapularis_Unknown_Drag_2016-12-01 | OL704193 |
| CAES | PV111-69_LII_Westport_CT_USA_Ixodes-scapularis_F_Drag_2019-04-09 | OL704194 |
| CAES | PV112-140_LII_Lyme_CT_USA_Ixodes-scapularis_F_Drag_2019-04-17 | OL704195 |
| CAES | PV113-364_LII_Westport_CT_USA_Ixodes-scapularis_F_Drag_2019-05-07 | OL704196 |
| CAES | PV114-375_LII_North-Branford_CT_USA_Ixodes-scapularis_F_Drag_2019-05-08 | OL704197 |
| Cornell | PV299-LI108_LII_Mill-Neck_NY_USA_Ixodes-scapularis_F_Drag_2018-10-25 | OL704210 |
| Cornell | PV300-LI131_LII_Mill-Neck_NY_USA_Ixodes-scapularis_M_Drag_2018-10-25 | OL704209 |
| Cornell | PV301-LI132_LII_Mill-Neck_NY_USA_Ixodes-scapularis_M_Drag_2018-10-25 | OL704208 |
| Cornell | PV302-LI171_LII_Mill-Neck_NY_USA_Ixodes-scapularis_F_Drag_2018-10-25 | OL704207 |
| Cornell | PV303-LI192_LII_Mill-Neck_NY_USA_Ixodes-scapularis_M_Drag_2018-10-25 | OL704206 |
| Cornell | PV305-LI212_LII_Mill-Neck_NY_USA_Ixodes-scapularis_M_Drag_2018-10-25 | OL704205 |
| Cornell | PV306-LI298_LII_Oyster-Bay_NY_USA_Ixodes-scapularis_F_Drag_2018-10-31 | OL704204 |
| Cornell | PV307-LI305_LII_Oyster-Bay_NY_USA_Ixodes-scapularis_F_Drag_2018-10-31 | OL704203 |
| Cornell | PV309-LI327_LII_Oyster-Bay_NY_USA_Ixodes-scapularis_M_Drag_2018-10-31 | OL704202 |
| Cornell | PV310-LI331_LII_Oyster-Bay_NY_USA_Ixodes-scapularis_M_Drag_2018-10-31 | OL704201 |
| Cornell | PV311-LI336_LII_Oyster-Bay_NY_USA_Ixodes-scapularis_M_Drag_2018-10-31 | OL704200 |
| Cornell | PV312-LI343_LII_Oyster-Bay_NY_USA_Ixodes-scapularis_M_Drag_2018-10-31 | OL704199 |
| Cornell | PV313-LI586_LII_Woodbury_NY_USA_Ixodes-scapularis_M_Drag_2018-12-04 | OL704198 |
| MHIR | PV083-242_LII_Cape-Elizabeth_ME_USA_Ixodes-scapularis_N_Drag_2016-06-13 | OL704212 |
| MHIR | PV084-612_LII_Wells_ME_USA_Ixodes-scapularis_F_Drag_2016-10-07 | OL704213 |
| MHIR | PV085-632_LII_Wells_ME_USA_Ixodes-scapularis_F_Drag_2016-10-07 | OL704214 |
| MHIR | PV086-687_LII_Midcoast_ME_USA_Ixodes-scapularis_F_Drag_2016-10-14 | OL704215 |
| MHIR | PV087-946_LII_Cape-Elizabeth_ME_USA_Ixodes-scapularis_M_Drag_2016-10-06 | OL704216 |
| MHIR | PV088-963_LII_Wells_ME_USA_Ixodes-scapularis_M_Drag_2016-11-07 | OL704217 |
| MHIR | PV089-1003_LII_Wells_ME_USA_Ixodes-scapularis_F_Drag_2016-11-07 | OL704218 |
| MHIR | PV090-1135_LII_Wells_ME_USA_Ixodes-scapularis_M_Drag_2016-11-09 | OL704219 |
| MHIR | PV091-1141_LII_Wells_ME_USA_Ixodes-scapularis_M_Drag_2016-11-09 | OL704220 |
| MHIR | PV093-1331_LII_Standish_ME_USA_Ixodes-scapularis_M_Drag_2016-11-17 | OL704222 |
| MHIR | PV094-1341_LII_Standish_ME_USA_Ixodes-scapularis_M_Drag_2016-11-17 | OL704223 |
| MHIR | PV096-CO18_LII_Cape-Elizabeth_ME_USA_Ixodes-scapularis_M_Drag_2017-11-15 | OL704224 |
| MHIR | PV098-CO57_LII_Cape-Elizabeth_ME_USA_Ixodes-scapularis_F_Drag_2017-11-15 | OL704226 |
| MHIR | PV102-CO166_LII_Cape-Elizabeth_ME_USA_Ixodes-scapularis_F_Drag_2017-11-15 | OL704230 |
| MHIR | PV103-CO527_LII_Cape-Elizabeth_ME_USA_Ixodes-scapularis_M_Drag_2017-11-15 | OL704231 |
| MHIR | PV105-CO570_LII_Cape-Elizabeth_ME_USA_Ixodes-scapularis_M_Drag_2017-11-07 | OL704233 |
| MHIR | PV106-CO574_LII_Cape-Elizabeth_ME_USA_Ixodes-scapularis_M_Drag_2017-11-07 | OL704234 |
| MHIR | PV109-CO452_LII_Wells_ME_USA_Ixodes-scapularis_F_Drag_2017-11-15 | OL704236 |
| MHIR | PV286-POW19-088_LII_Thomaston_ME_USA_Ixodes-scapularis_M_Drag_2019-05-22 | OL704237 |
| MHIR | PV287-WB19-061_LII_Wells_ME_USA_Ixodes-scapularis_N_Bird_2019-06-18 | OL704238 |
| MHIR | PV292-PF3.019_LII_Wells_ME_USA_Ixodes-scapularis_M_Drag_2018-06-07 | OL704239 |
| MHIR | PV293-PF3.020_LII_Wells_ME_USA_Ixodes-scapularis_M_Drag_2018-06-07 | OL704240 |
| MHIR | PV297-PF1.021_LII_Wells_ME_USA_Ixodes-scapularis_M_Drag_2018-06-19 | OL704241 |
| MHIR | PV298-PF6.005_LII_Wells_ME_USA_Ixodes-scapularis_M_Drag_2018-06-14 | OL704242 |
| MHIR | PV081.2-H11_LII_Cape-Elizabeth_ME_USA_Ixodes-scapularis_M_Drag_2016-04-08 | OL704211 |
| MHIR | PV092.2-1217_LII_Cape-Elizabeth_ME_USA_Ixodes-scapularis_F_Drag_2016-11-18 | OL704221 |
| MHIR | PV097.2-CO46_LII_Cape-Elizabeth_ME_USA_Ixodes-scapularis_F_Drag_2017-11-15 | OL704225 |
| MHIR | PV099.2-CO68_LII_Cape-Elizabeth_ME_USA_Ixodes-scapularis_F_Drag_2017-11-15 | OL704227 |
| MHIR | PV100.2-CO95_LII_Cape-Elizabeth_ME_USA_Ixodes-scapularis_M_Drag_2017-11-15 | OL704228 |
| MHIR | PV101.2-CO110_LII_Cape-Elizabeth_ME_USA_Ixodes-scapularis_F_Drag_2017-11-15 | OL704229 |
| MHIR | PV104.2-CO542_LII_Cape-Elizabeth_ME_USA_Ixodes-scapularis_F_Drag_2017-11-15 | OL704232 |
| MHIR | PV108.2-CO239_LII_Wells_ME_USA_Ixodes-scapularis_M_Drag_2017-11-15 | OL704235 |
| NYSDOH | PV116-18079-121_LII_Brewster_NY_USA_Ixodes-scapularis_M_Drag_2018-10-24 | OL704243 |
| NYSDOH | PV117-18087-108_LII_Montebello_NY_USA_Ixodes-scapularis_F_Drag_2018-10-26 | OL704244 |
| NYSDOH | PV118-18087-113_LII_Montebello_NY_USA_Ixodes-scapularis_M_Drag_2018-10-26 | OL704245 |
| NYSDOH | PV119-18119-217_LII_Mohansic_NY_USA_Ixodes-scapularis_M_Drag_2018-10-22 | OL704246 |
| NYSDOH | PV120-18119-218_LII_Mohansic_NY_USA_Ixodes-scapularis_M_Drag_2018-10-22 | OL704247 |
| NYSDOH | PV121-18119-219_LII_Mohansic_NY_USA_Ixodes-scapularis_M_Drag_2018-10-22 | OL704248 |
| NYSDOH | PV122-18119-228_LII_Armonk_NY_USA_Ixodes-scapularis_M_Drag_2018-10-31 | OL704249 |
| NYSDOH | PV123-17091-200_LII_Saratoga-Springs_NY_USA_Ixodes-scapularis_N_Drag_2017-07-20 | OL704250 |
| NYSDOH | PV124-17119-057_LII_Armonk_NY_USA_Ixodes-scapularis_N_Drag_2017-06-13 | OL704251 |
| NYSDOH | PV125-17119-059_LII_Armonk_NY_USA_Ixodes-scapularis_N_Drag_2017-06-13 | OL704252 |
| NYSDOH | PV127-17091-230_LII_Stillwater_NY_USA_Ixodes-scapularis_N_Drag_2017-07-21 | OL704253 |
| NYSDOH | PV128-17091-144_LII_Malta_NY_USA_Ixodes-scapularis_F_Drag_2017-07-05 | OL704254 |
| NYSDOH | PV129-18095-057_LII_Blenheim_NY_USA_Ixodes-scapularis_F_Drag_2018-06-15 | OL704255 |
| NYSDOH | PV130-18021-061_LII_Clermont_NY_USA_Ixodes-scapularis_F_Drag_2018-05-24 | OL704256 |
| NYSDOH | PV131-18021-063_LII_Clermont_NY_USA_Ixodes-scapularis_F_Drag_2018-05-24 | OL704257 |
| NYSDOH | PV132-17119-2-009_LII_Redding_CT_USA_Ixodes-scapularis_M_Deer_2017-10-28 | OL704258 |
| NYSDOH | PV133-17021-480_LII_Clermont_NY_USA_Ixodes-scapularis_F_Drag_2017-11-01 | OL704259 |
| NYSDOH | PV134-17091-236_LII_Greenfield_NY_USA_Ixodes-scapularis_N_Drag_2017-07-20 | OL704260 |
| NYSDOH | PV135-17021-482_LII_Clermont_NY_USA_Ixodes-scapularis_F_Drag_2017-11-01 | OL704261 |
| NYSDOH | PV136-18039-055_LII_Cairo_NY_USA_Ixodes-scapularis_L_Drag_2018-05-31 | OL704262 |
| NYSDOH | PV137-16021-172_LII_Greenport_NY_USA_Ixodes-scapularis_M_Drag_2016-10-21 | OL704263 |
| NYSDOH | PV138-15-S22-01_LII_Shelter-Island_NY_USA_Ixodes-scapularis_F_Deer_2016-01-07 | OL704264 |
| NYSDOH | PV139-15071-117_LII_Wallkill_NY_USA_Ixodes-scapularis_M_Drag_2015-10-21 | OL704265 |
| NYSDOH | PV141-17091-11-006_LII_Stillwater_NY_USA_Ixodes-scapularis_M_Deer_2017-11-18 | OL704266 |
| NYSDOH | PV142-17091-23-001_LII_Saratoga_NY_USA_Ixodes-scapularis_M_Deer_2017-11-18 | OL704267 |
| NYSDOH | PV143-17119-131_LII_Armonk_NY_USA_Ixodes-scapularis_M_Drag_2017-10-25 | OL704268 |
| NYSDOH | PV144-17091-27-001_LII_Clifton-Park_NY_USA_Ixodes-scapularis_M_Deer_2017-11-19 | OL704269 |
| NYSDOH | PV145-17091-11-002_LII_Stillwater_NY_USA_Ixodes-scapularis_M_Deer_2017-11-18 | OL704270 |
| NYSDOH | PV146-17027-241_LII_Holmes_NY_USA_Ixodes-scapularis_M_Drag_2017-10-23 | OL704271 |
| NYSDOH | PV147-17119-27-001_LII_Somers_NY_USA_Ixodes-scapularis_M_Deer_2017-11-08 | OL704272 |
| NYSDOH | PV148-17079-117_LII_Brewster_NY_USA_Ixodes-scapularis_M_Drag_2017-10-16 | OL704273 |
| NYSDOH | PV149-17103-304_LII_Islip_NY_USA_Ixodes-scapularis_M_Drag_2017-05-08 | OL704274 |
| NYSDOH | PV150-17103-254_LII_Huntington_NY_USA_Ixodes-scapularis_F_Drag_2017-05-12 | OL704275 |
| NYSDOH | PV151-17119-18-001_LII_Pound-Ridge_NY_USA_Ixodes-scapularis_M_Deer_2017-10-28 | OL704276 |
| NYSDOH | PV152-12021-250_LII_Clermont_NY_USA_Ixodes-scapularis_M_Drag_2012-11-16 | OL704277 |
| NYSDOH | PV153-14CD06-001_LII_Ancram_NY_USA_Ixodes-scapularis_M_Deer_2014-11-15 | OL704278 |
| NYSDOH | PV154-14CD13-002_LII_Copake_NY_USA_Ixodes-scapularis_M_Deer_2014-11-15 | OL704279 |
| NYSDOH | PV155-14CD34-002_LII_North-Hoosick_NY_USA_Ixodes-scapularis_M_Deer_2014-11-16 | OL704280 |
| NYSDOH | PV156-14071-112_LII_Wallkill_NY_USA_Ixodes-scapularis_M_Drag_2014-10-28 | OL704281 |
| NYSDOH | PV157-14071-113_LII_Wallkill_NY_USA_Ixodes-scapularis_M_Drag_2014-10-28 | OL704282 |
| NYSDOH | PV158-14071-118_LII_Wallkill_NY_USA_Ixodes-scapularis_F_Drag_2014-10-28 | OL704283 |
| NYSDOH | PV159-14071-119_LII_Wallkill_NY_USA_Ixodes-scapularis_F_Drag_2014-10-28 | OL704284 |
| NYSDOH | PV160-14103-145_LII_Islip_NY_USA_Ixodes-scapularis_M_Drag_2014-11-10 | OL704285 |
| NYSDOH | PV161-15CD01-001_LII_East-Taghkanic_NY_USA_Ixodes-scapularis_M_Deer_2015-11-21 | OL704286 |
| NYSDOH | PV162-15CD15-001_LII_Ancramdale_NY_USA_Ixodes-scapularis_M_Deer_2015-11-21 | OL704287 |
| NYSDOH | PV163-15CD31-001_LII_Copake_NY_USA_Ixodes-scapularis_M_Deer_2015-11-22 | OL704288 |
| NYSDOH | PV164-15CD59-002_LII_Poestenkill_NY_USA_Ixodes-scapularis_M_Deer_2015-11-22 | OL704289 |
| NYSDOH | PV165-12021-240_LII_Clermont_NY_USA_Ixodes-scapularis_F_Drag_2012-11-16 | OL704290 |
| NYSDOH | PV166-18103-2025_LII_Huntington_NY_USA_Ixodes-scapularis_F_Drag_2018-11-19 | OL704291 |
| NYSDOH | PV167-18103-1656_LII_East-Hampton_NY_USA_Ixodes-scapularis_F_Drag_2018-10-23 | OL704292 |
| NYSDOH | PV168-18103-1536_LII_Islip_NY_USA_Ixodes-scapularis_F_Drag_2018-10-19 | OL704293 |
| NYSDOH | PV169-19001-410_LII_Guilderland_NY_USA_Ixodes-scapularis_F_Drag_2019-04-17 | OL704294 |
| NYSDOH | PV170-19001-415_LII_Guilderland_NY_USA_Ixodes-scapularis_F_Drag_2019-04-17 | OL704295 |
| NYSDOH | PV171-19001-427_LII_Guilderland_NY_USA_Ixodes-scapularis_F_Drag_2019-04-17 | OL704296 |
| NYSDOH | PV172-19001-589_LII_Guilderland_NY_USA_Ixodes-scapularis_F_Drag_2019-05-22 | OL704297 |
| NYSDOH | PV173-19001-591_LII_Guilderland_NY_USA_Ixodes-scapularis_F_Drag_2019-05-22 | OL704298 |
| NYSDOH | PV174-19001-601_LII_Guilderland_NY_USA_Ixodes-scapularis_M_Drag_2019-05-22 | OL704299 |
| NYSDOH | PV175-19021-063_LII_Chatham_NY_USA_Ixodes-scapularis_F_Drag_2019-05-07 | OL704300 |
| NYSDOH | PV176-19039-010_LII_Greenville_NY_USA_Ixodes-scapularis_M_Drag_2019-05-08 | OL704301 |
| NYSDOH | PV177-19091-035_LII_Wilton_NY_USA_Ixodes-scapularis_F_Drag_2019-04-22 | OL704302 |
| NYSDOH | PV178-19091-043_LII_Wilton_NY_USA_Ixodes-scapularis_M_Drag_2019-04-22 | OL704303 |
| NYSDOH | PV179-19091-084_LII_Saratoga-Springs_NY_USA_Ixodes-scapularis_F_Drag_2019-04-29 | OL704304 |
| NYSDOH | PV180-19091-164_LII_Wilton_NY_USA_Ixodes-scapularis_F_Drag_2019-05-15 | OL704305 |
| NYSDOH | PV181-19091-165_LII_Wilton_NY_USA_Ixodes-scapularis_F_Drag_2019-05-15 | OL704306 |
| NYSDOH | PV182-19093-125_LII_Glenville_NY_USA_Ixodes-scapularis_F_Drag_2019-05-16 | OL704307 |
| NYSDOH | PV183-19093-142_LII_Glenville_NY_USA_Ixodes-scapularis_M_Drag_2019-05-16 | OL704308 |
| NYSDOH | PV185-19093-242_LII_Glenville_NY_USA_Ixodes-scapularis_M_Drag_2019-05-22 | OL704309 |
| NYSDOH | PV186-19115-021_LII_Jackson_NY_USA_Ixodes-scapularis_F_Drag_2019-05-06 | OL704310 |
| NYSDOH | PV187-16CD17_LII_Copake_NY_USA_Ixodes-scapularis_M_Deer_2016-11-19 | OL704311 |
| NYSDOH | PV188-16021-171_LII_Greenport_NY_USA_Ixodes-scapularis_F_Drag_2016-10-21 | OL704312 |
| NYSDOH | PV189-18091-387_LII_Charlton_NY_USA_Ixodes-scapularis_M_Drag_2018-11-08 | OL704313 |
| NYSDOH | PV190-18091-16-005_LII_Stillwater_NY_USA_Ixodes-scapularis_M_Deer_2018-11-17 | OL704314 |
| NYSDOH | PV191-18091-26-001_LII_Schaghticoke_NY_USA_Ixodes-scapularis_M_Deer_2018-11-18 | OL704315 |
| NYSDOH | PV192-18091-11-003_LII_Saratoga_NY_USA_Ixodes-scapularis_M_Deer_2018-11-17 | OL704316 |
| NYSDOH | PV193-18091-328_LII_Milton_NY_USA_Ixodes-scapularis_M_Drag_2018-11-07 | OL704317 |
| NYSDOH | PV194-18089-048_LII_Hermon_NY_USA_Ixodes-scapularis_F_Drag_2018-10-14 | OL704318 |
| NYSDOH | PV195-18089-049_LII_Hermon_NY_USA_Ixodes-scapularis_F_Drag_2018-10-14 | OL704319 |
| NYSDOH | PV196-15-S13-03_LII_Shelter-Island_NY_USA_Ixodes-scapularis_M_Deer_2016-01-13 | OL704320 |
| NYSDOH | PV197-15-S26-01_LII_Shelter-Island_NY_USA_Ixodes-scapularis_M_Deer_2016-01-20 | OL704321 |
| NYSDOH | PV200-15CD63-001_LII_Nassau_NY_USA_Ixodes-scapularis_M_Deer_2015-11-22 | OL704322 |
| NYSDOH | PV201-16021-231_LII_Clermont_NY_USA_Ixodes-scapularis_M_Drag_2016-10-25 | OL704323 |
| NYSDOH | PV202-17091-180_LII_Saratoga-Springs_NY_USA_Ixodes-scapularis_N_Drag_2017-07-20 | OL704324 |
| NYSDOH | PV203-17021-227_LII_Greenport_NY_USA_Ixodes-scapularis_N_Drag_2017-05-24 | OL704325 |
| NYSDOH | PV204-19103-1357_LII_Southampton_NY_USA_Ixodes-scapularis_N_Drag_2019-06-17 | OL704326 |
| NYSDOH | PV205-19119-078_LII_Armonk_NY_USA_Ixodes-scapularis_N_Drag_2019-05-31 | OL704327 |
| NYSDOH | PV206-19103-01-001_LII_Shelter-Island_NY_USA_Ixodes-scapularis_M_Deer_2019-01-07 | OL704328 |
| NYSDOH | PV207-19103-14-001_LII_Shelter-Island_NY_USA_Ixodes-scapularis_M_Deer_2019-01-07 | OL704329 |
| NYSDOH | PV208-19103-27-001_LII_Shelter-Island_NY_USA_Ixodes-scapularis_M_Deer_2019-01-08 | OL704330 |
| NYSDOH | PV209-19103-36-003_LII_Shelter-Island_NY_USA_Ixodes-scapularis_F_Deer_2019-01-08 | OL704331 |
| NYSDOH | PV210-19103-37-002_LII_Shelter-Island_NY_USA_Ixodes-scapularis_M_Deer_2019-01-08 | OL704332 |
| NYSDOH | PV211-19103-37-004_LII_Shelter-Island_NY_USA_Ixodes-scapularis_F_Deer_2019-01-08 | OL704333 |
| NYSDOH | PV212-19103-41-001_LII_Shelter-Island_NY_USA_Ixodes-scapularis_M_Deer_2019-01-08 | OL704334 |
| NYSDOH | PV213-19103-43-001_LII_Shelter-Island_NY_USA_Ixodes-scapularis_F_Deer_2019-01-09 | OL704335 |
| NYSDOH | PV215-19091-227_LII_Clifton-Park_NY_USA_Ixodes-scapularis_F_Drag_2019-05-31 | OL704336 |
| NYSDOH | PV216-19091-231_LII_Clifton-Park_NY_USA_Ixodes-scapularis_M_Drag_2019-05-31 | OL704337 |
| NYSDOH | PV217-19095-165_LII_North-Blenheim_NY_USA_Ixodes-scapularis_M_Drag_2019-05-30 | OL704338 |
| NYSDOH | PV218-13CD01-002_LII_Copake_NY_USA_Ixodes-scapularis_M_Deer_2013-11-16 | OL704339 |
| NYSDOH | PV222-14021-346_LII_Clermont_NY_USA_Ixodes-scapularis_F_Drag_2014-11-25 | OL704340 |
| NYSDOH | PV223-14021-358_LII_Clermont_NY_USA_Ixodes-scapularis_M_Drag_2014-11-25 | OL704341 |
| NYSDOH | PV224-14021-472_LII_Chatham_NY_USA_Ixodes-scapularis_M_Drag_2014-11-25 | OL704342 |
| NYSDOH | PV227-14087-111_LII_Montebello_NY_USA_Ixodes-scapularis_M_Drag_2014-11-04 | OL704343 |
| NYSDOH | PV228-14105-140_LII_Mamakating_NY_USA_Ixodes-scapularis_M_Drag_2014-10-28 | OL704344 |
| NYSDOH | PV229-17091-15-003_LII_Stillwater_NY_USA_Ixodes-scapularis_M_Deer_2017-11-18 | OL704345 |
| NYSDOH | PV230-17105-109_LII_Mamakating_NY_USA_Ixodes-scapularis_F_Drag_2017-10-27 | OL704346 |
| NYSDOH | PV231-17119-29-002_LII_Pound-Ridge_NY_USA_Ixodes-scapularis_M_Deer_2017-11-13 | OL704347 |
| NYSDOH | PV232-18103-167-010_LII_Shelter-Island_NY_USA_Ixodes-scapularis_F_Deer_2018-11-01 | OL704348 |
| NYSDOH | PV233-18103-168-007_LII_Shelter-Island_NY_USA_Ixodes-scapularis_Unknown_Deer_2018-11-01 | OL704349 |
| NYSDOH | PV234-18071-054_LII_Wallkill_NY_USA_Ixodes-scapularis_N_Drag_2018-11-06 | OL704350 |
| NYSDOH | PV235-18079-053_LII_Brewster_NY_USA_Ixodes-scapularis_N_Drag_2018-05-31 | OL704351 |
| NYSDOH | PV236-18087-055_LII_Montebello_NY_USA_Ixodes-scapularis_N_Drag_2018-06-18 | OL704352 |
| NYSDOH | PV237-18091-128_LII_Saratoga-Springs_NY_USA_Ixodes-scapularis_M_Drag_2018-06-27 | OL704353 |
| NYSDOH | PV239-18103-058_LII_Smithtown_NY_USA_Ixodes-scapularis_F_Drag_2018-02-27 | OL704354 |
| NYSDOH | PV240-18103-1315_LII_Huntington_NY_USA_Ixodes-scapularis_N_Drag_2018-06-07 | OL704355 |
| NYSDOH | PV241-18103-1419_LII_Smithtown_NY_USA_Ixodes-scapularis_N_Drag_2018-06-06 | OL704356 |
| NYSDOH | PV242-18103-281_LII_East-Hampton_NY_USA_Ixodes-scapularis_N_Drag_2018-06-19 | OL704357 |
| NYSDOH | PV243-17021-304_LII_Greenport_NY_USA_Ixodes-scapularis_F_Drag_2017-11-01 | OL704358 |
| NYSDOH | PV244-17021-341_LII_Taghkanic_NY_USA_Ixodes-scapularis_F_Drag_2017-11-01 | OL704359 |
| NYSDOH | PV245-17091-11-001_LII_Stillwater_NY_USA_Ixodes-scapularis_M_Deer_2017-11-18 | OL704360 |
| NYSDOH | PV246-17091-23-002_LII_Saratoga_NY_USA_Ixodes-scapularis_M_Deer_2017-11-18 | OL704361 |
| NYSDOH | PV247-17091-24-003_LII_Ballston-Spa_NY_USA_Ixodes-scapularis_M_Deer_2017-11-18 | OL704362 |
| NYSDOH | PV248-17091-27-004_LII_Clifton-Park_NY_USA_Ixodes-scapularis_M_Deer_2017-11-19 | OL704363 |
| NYSDOH | PV249-17091-391_LII_Wilton_NY_USA_Ixodes-scapularis_F_Drag_2017-10-27 | OL704364 |
| NYSDOH | PV250-17091-457_LII_Saratoga-Springs_NY_USA_Ixodes-scapularis_F_Drag_2017-11-09 | OL704365 |
| NYSDOH | PV251-17091-474_LII_Greenfield_NY_USA_Ixodes-scapularis_F_Drag_2017-11-09 | OL704366 |
| NYSDOH | PV252-17091-500_LII_Northumberland_NY_USA_Ixodes-scapularis_M_Drag_2017-11-15 | OL704367 |
| NYSDOH | PV253-17091-536_LII_Stillwater_NY_USA_Ixodes-scapularis_M_Drag_2017-11-15 | OL704368 |
| NYSDOH | PV254-17091-654_LII_Malta_NY_USA_Ixodes-scapularis_F_Drag_2017-11-09 | OL704369 |
| NYSDOH | PV255-17091-658_LII_Malta_NY_USA_Ixodes-scapularis_M_Drag_2017-11-09 | OL704370 |
| NYSDOH | PV256-17091-660_LII_Malta_NY_USA_Ixodes-scapularis_M_Drag_2017-11-09 | OL704371 |
| NYSDOH | PV257-17103-1037_LII_Islip_NY_USA_Ixodes-scapularis_F_Drag_2017-11-21 | OL704372 |
| NYSDOH | PV258-17103-1045_LII_Huntington_NY_USA_Ixodes-scapularis_M_Drag_2017-10-27 | OL704373 |
| NYSDOH | PV259-17103-973_LII_East-Hampton_NY_USA_Ixodes-scapularis_M_Drag_2017-11-15 | OL704374 |
| NYSDOH | PV260-17115-194_LII_Greenwich_NY_USA_Ixodes-scapularis_M_Drag_2017-10-27 | OL704375 |
| NYSDOH | PV261-19001-855_LII_Guilderland_NY_USA_Ixodes-scapularis_F_Drag_2019-10-15 | OL704376 |
| NYSDOH | PV262-19001-866_LII_Guilderland_NY_USA_Ixodes-scapularis_F_Drag_2019-10-15 | OL704377 |
| NYSDOH | PV263-19001-899_LII_Guilderland_NY_USA_Ixodes-scapularis_M_Drag_2019-10-15 | OL704378 |
| NYSDOH | PV264-19021-783_LII_Copake_NY_USA_Ixodes-scapularis_F_Drag_2019-11-06 | OL704379 |
| NYSDOH | PV265-19021-794_LII_Copake_NY_USA_Ixodes-scapularis_F_Drag_2019-11-06 | OL704380 |
| NYSDOH | PV266-19039-373_LII_Windham_NY_USA_Ixodes-scapularis_M_Drag_2019-10-25 | OL704381 |
| NYSDOH | PV267-19039-497_LII_Cairo_NY_USA_Ixodes-scapularis_M_Drag_2019-10-25 | OL704382 |
| NYSDOH | PV268-19083-510_LII_Schodack_NY_USA_Ixodes-scapularis_F_Drag_2019-11-05 | OL704383 |
| NYSDOH | PV269-19083-516_LII_Schodack_NY_USA_Ixodes-scapularis_F_Drag_2019-11-05 | OL704384 |
| NYSDOH | PV270-19091-625_LII_Malta_NY_USA_Ixodes-scapularis_F_Drag_2019-10-10 | OL704385 |
| NYSDOH | PV271-19093-494_LII_Glenville_NY_USA_Ixodes-scapularis_F_Drag_2019-11-04 | OL704386 |
| NYSDOH | PV272-19065-058_LI_Verona_NY_USA_Ixodes-scapularis_M_Drag_2019-11-04 | OL704387 |
| NYSDOH | PV273-19065-059_LI_Verona_NY_USA_Ixodes-scapularis_F_Drag_2019-11-04 | OL704388 |
| NYSDOH | PV274-19021-02-002_LII_Ancram_NY_USA_Ixodes-scapularis_M_Deer_2019-11-16 | OL704389 |
| NYSDOH | PV275-19021-03-002_LII_North-East_NY_USA_Ixodes-scapularis_M_Deer_2019-11-16 | OL704390 |
| NYSDOH | PV276-19021-09-001_LII_Taghkanic_NY_USA_Ixodes-scapularis_M_Deer_2019-11-16 | OL704391 |
| NYSDOH | PV277-19091-04-001_LII_Ballston_NY_USA_Ixodes-scapularis_M_Deer_2019-11-16 | OL704392 |
| NYSDOH | PV278-19091-08-002_LII_Stillwater_NY_USA_Ixodes-scapularis_M_Deer_2019-11-16 | OL704393 |
| NYSDOH | PV279-19091-11-001_LII_Greenwich_NY_USA_Ixodes-scapularis_M_Deer_2019-11-16 | OL704394 |

**Table S4:**
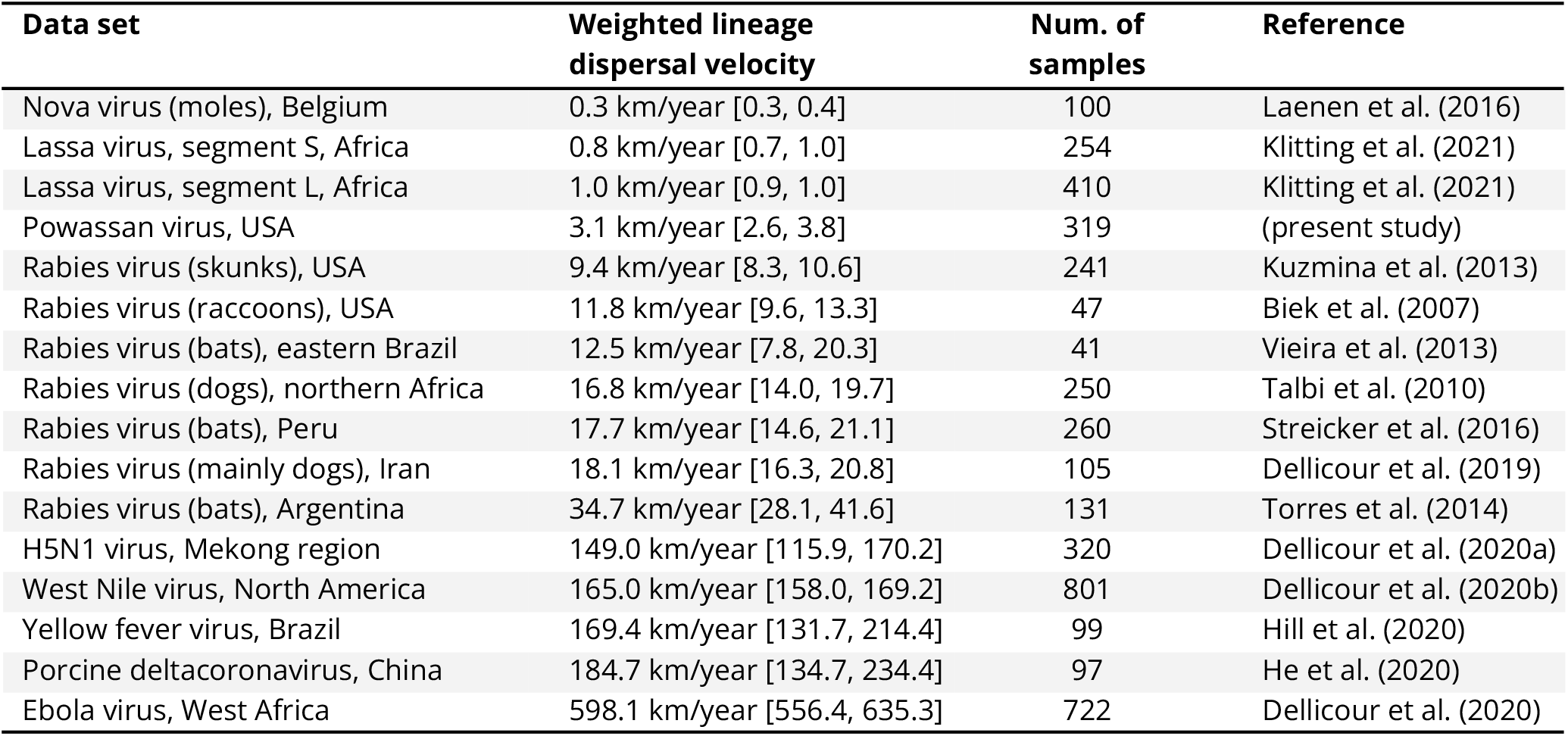
Comparison of lineage dispersal velocities estimated for different data sets of viral genomes. The table comes from the study of Klitting et al. (2021) (79) and has been completed with the estimate obtained from the continuous phylogeographic reconstruction of Powassan virusOWV lineages in the northeastern United States. For each data set, we report both the posterior median estimate and the 95% HPD interval.

**Table S5:** Impact of several environmental factors on the dispersal velocity of Powassan virus lineages. The results are based on 1,000 posterior trees obtained by spatially-explicit phylogeographic inference. “C” and “R” indicate if the considered environmental raster was considered as a conductance (“C”) or resistance factor (“R”), and *k* is the rescaling parameter used to transform the initial raster (see the text for further details). For regression coefficients and *Q* values we report both the median estimate and the 95% HPD interval. The Bayes factor (BF) supports are only reported when p(*Q* > 0) is at least 90%. Following Kass & Raftery (1995), we consider a BF value >20 as strong support for a significant correlation between the environmental distances and dispersal durations (in bold).

| Environmental factor | Tendency of viral lineages to avoid circulating within specific environmental conditions | Tendency of viral lineages to preferentially circulate within specific environmental conditions |
| --- | --- | --- |
| Elevation | >99 | 0.0 |
| Forest areas | 9.6 | 0.1 |
| Croplands | 0.2 | 5.9 |
| Urban areas | 0.4 | 2.8 |
| Annual mean Temperature | 0.1 | 13.3 |
| Annual precipitation | 8.7 | 0.1 |

**Table S6:**
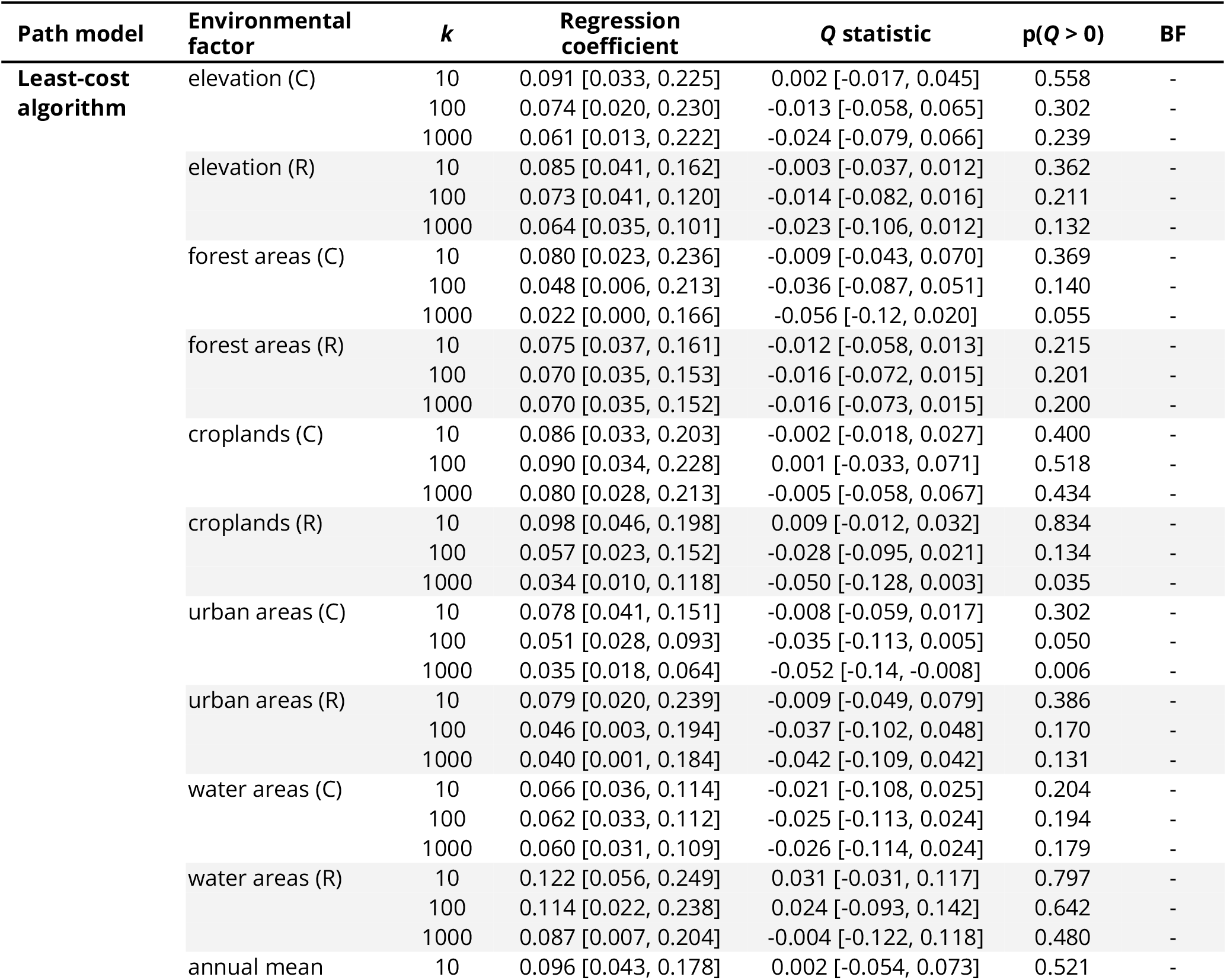
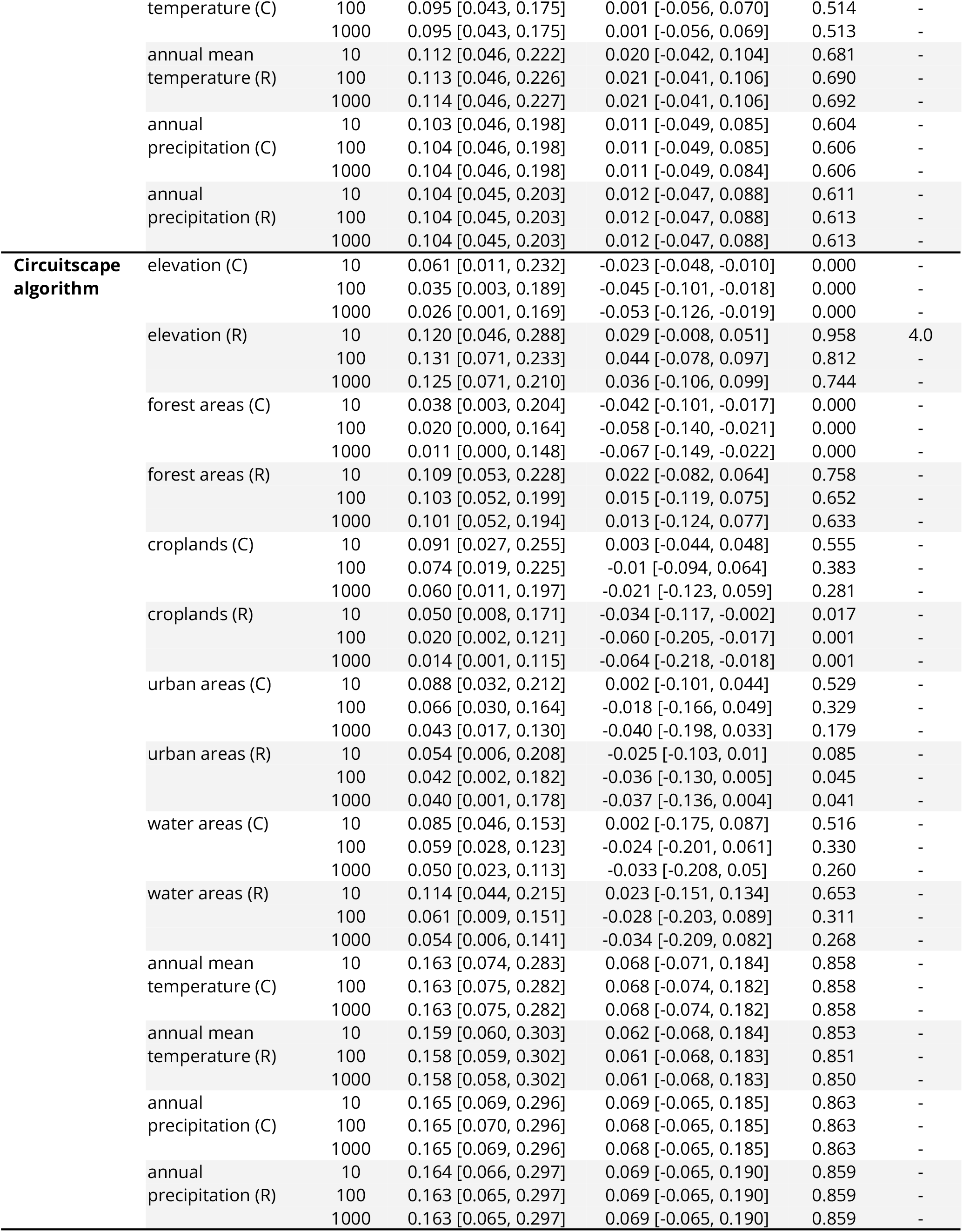
Impact of several environmental factors on the dispersal location of Powassan virus lineages. We report Bayes factor (BF) support for the association between environmental values and tree node locations. The results are based on 1,000 posterior trees obtained by spatially-explicit phylogeographic inference. Following Kass & Raftery (1995), we consider a BF value >20 as strong support for a significant correlation between the environmental distances and dispersal durations (in bold). “ENM” refers to ecological niche modeling.

## Notes

### Summary of Updates

Included details on priors used in the BETS analysis and provided trees, "xml" files, and log files for phylogeographic analyses.

